# Layer 5 lateral entorhinal cortex neurons encode updating of object-place associations

**DOI:** 10.1101/2025.08.29.673029

**Authors:** PJ Banks, GRI Barker, CA Booth, L Kinnavane, EC Warburton, ZI Bashir

## Abstract

Associative recognition memory is essential for everyday life, forming cognitive representations of and recalling relationships between things we encounter and their environments. Object-place associations are represented in the lateral entorhinal and medial prefrontal cortices, however the identity of neurons in which these associations are formed, the cellular mechanisms supporting them, and how these representations react to change are not understood. Here we labelled associative recognition memory engrams, finding that engram neuron reactivation by memory recall correlated with behavioural performance only in the dorsolateral subregion of entorhinal cortex, where reactivation was overrepresented in layer 5/6. Electrophysiology from e*x vivo* slices prepared directly following memory recall revealed increased excitability in layer 5 lateral entorhinal cortex engram cells, which only occurred when engram-specific objects were reconfigured. These data identify deep layer lateral entorhinal cortex neurons as key loci of object-place associations and proposes a plastic mechanism by which pre-existing neural representations are updated.

**Significance statement:** Associative recognition memory is key to our normal everyday lives, but how these memories are updated and which neurons support these memories remain poorly understood. We identify a population of neurons in the deep-layers of lateral entorhinal cortex which are activated by memory encoding and whose reactivation is correlated with, and necessary for, memory retrieval, thus identifying them as engram neurons. These entorhinal neurons demonstrate increased firing following presentation of familiar cues in a novel configuration, revealing a mechanism for integrating new information into existing memories. By identifying a specific cell type and plasticity process underlying memory updating, this work advances understanding of how flexible object-place representations are maintained in the brain.

## Introduction

Associative recognition memory, remembrance for objects and their associated locations and contexts, is fundamental to everyday decision making and behaviour. These memories depend on a highly interconnected brain network including the hippocampus, thalamus, perirhinal cortex (PRH), medial prefrontal cortex (mPFC) and medial and lateral entorhinal cortex (MEC, LEC; Aggleton and Nelson, 2020). Of these cortical regions, PRH has a well-established role encoding object identity and novelty, whilst mPFC performs integration of object identity, location and time (Ennaceur et al., 1997; Barker et al., 2007). The role of LEC, however, is less clear.

Classical models (Eichenbaum et al., 2007) propose that LEC is part of a processing stream representing object features and identity, which converges with spatial information from MEC to form object-context representations in the hippocampus. Emerging evidence however suggests that LEC neurons can represent dimensions not only of object identity but also of context, time and space (Deshmukh and Knierim, 2011; Deshmukh et al., 2012; Keene et al., 2016; Tsao et al., 2018; Wang et al., 2018; Nilssen et al., 2019; Wang et al., 2020; Wang et al., 2024). Indeed, lesion or inactivation of LEC impairs object-in-place (OiP) and object-in-context (OiC) memory whilst sparing simple novel object recognition (Van Cauter et al., 2013; Wilson et al., 2013; Chao et al., 2016; Kinnavane et al., 2025; though see: Vandrey et al., 2020).

Which neurons in these association cortices are prominently active during OiP memory and the plastic changes which may support memory is unclear; both superficial and deep LEC neurons have been shown to encode multi-dimensional object representations (Deshmukh and Knierim, 2011; Deshmukh et al., 2012; Keene et al., 2016). Optogenetic inhibition (Kinnavane et al., 2025) of mPFC→LEC synapses, which innervate deep layers of LEC (Kinnavane and Banks, 2022; Kinnavane et al., 2025), impairs both OiP and OiC memory whilst LEC layer 2a fan cells are crucial for associative learning (Lee et al., 2021) and object-context-place discrimination (Vandrey et al., 2020). For mPFC, immediate-early gene studies have shown activation of mPFC by associative recognition memory tasks, but have not examined laminar activation in detail (Barbosa et al., 2013), and layer specific *in vivo* recordings are lacking.

In this study, to identify key cell populations involved in associative memory processing, we used *Fos^2A-iCreERT2^* (TRAP2; Allen et al., 2017) mice to label putative OiP memory engram cells, focussing on association cortices. OiP produced robust TRAP2 labelling at memory encoding and Fos activation following retrieval. TRAP2 and Fos double-positive neurons (engram cells which had been reactivated by retrieval) were significantly more numerous in OiP mice compared to controls, however this reactivation correlated with memory performance only in dorsal subregions of LEC. Chemogenetic inactivation showed LEC TRAP2 cells to be necessary for expression of OiP memory recall. In *ex vivo* LEC brain slices prepared from mice which had recently performed memory retrieval, layer 5 (L5) pyramidal TRAP2 neurons showed increased excitability and firing compared to unlabelled neurons; these changes were not found in other cortical engram populations and only occurred when objects were presented in a reconfigured arrangement. These data suggest a key role for LEC L5 neurons in forming object-place associations and demonstrate an engram cell specific form of plasticity related to retrieval and integration of new information into existing object-place representations.

## Methods

### Animals

All animal procedures were conducted in accordance with the United Kingdom Animals (Scientific Procedures) Act (1986) under project licence number PP7058522 and were approved by the University of Bristol Ethical Review Committee. The following strains of mice were used: TRAP2 (Fos^2A-iCreERT2^): Fos^tm2.1(icre/ERT2)Luo^/J, JAX stock 030323 generated by L.Luo (Allen et al., 2017); Ai14 (: B6.Cg-Gt(ROSA)26Sor^tm14(CAG-tdTomato)Hze^/J, JAX stock 007914 generated by H.Zeng (Madisen et al., 2010)); hM4Di-DREADD (R26-LSL-Gi-DREADD): B6.129-Gt(ROSA)26Sor^tm1(CAG-CHRM4*,-mCitrine)Ute^/J, JAX stock 026219 generated by B.Roth and U.Hochgeschwender (Zhu et al., 2016). Mice were maintained on a reverse 12 h:12 h light:dark cycle (lights on 20:00-08:00) at constant temperature and humidity with *ad libitum* food and water. Home cage enrichment was present in the form of a cylindrical cardboard tube and a cellulose half-dome. Both male and female mice (sex determined by visual inspection of gonads by the animal breeding unit) were used at ≥10 weeks old. Mice were group housed in cages of 2-4 except for experiments involving cannulated hM4Di-DREADD mice which were single housed to prevent chewing of cannulae between cage-mates – for this reason, only male mice were used for these experiments due to increased tolerance of single housing(Davies, 2025).

### Surgery

Implantation of bilateral cannulae into LEC was performed under isoflurane general anaesthesia (3% induction, 2-3% maintenance) and secured in a stereotaxic frame using a mouse anaesthesia head holder and non-rupture ear bars (World Precision Instruments; WPI). Ophthalmic ointment was applied to the eyes to prevent drying, the scalp was sterilised with 4% chlorhexidine, and lidocaine (2%, 0.05 ml) was injected subcutaneously to provide local anaesthesia. A scalpel incision was made along the midline of the scalp and craniotomies were performed with a microdrill (WPI) and 0.5 mm burrs (Fine Science Tools). Guide cannula (26G, P1Technologies, Cat#C315GAS-5/SPC) were implanted into LEC (AP −3.0mm, ML ±4.2mm, DV - 3.2mm relative to bregma) and secured with gentamycin bone cement (DePuy international). The scalp incision was closed with surgical sutures and wound powder applied to the site. Mice were given 0.5 ml subcutaneous glucose saline for hydration. Analgesia was provided via: 1.5 µg intramuscular buprenorphine given at the end of surgery and 0.15 mg meloxicam given orally in a palatable solution of condensed milk 24 h prior- and 24 and 48 h post-surgery. Mice were allowed to recover for 2 weeks before undergoing behavioural procedures.

### Behavioural Procedures

#### Habituation

Mice were habituated to the entire experimental regimen except objects for at least 6 days prior to activity-dependent labelling. All translocation of mice between cages and behavioural apparatus was done using cardboard enrichment tubes (Davies et al., 2022). Mice were first transported from the home room to the behavioural room within 2 hours of dark phase onset where they would remain for a similar duration and timing to activity-dependent labelling day. Cage mates were split into individual experimental cages which had the same dimensions, bedding and enrichment as their home cages, and placed in a designated position in a storage rack surrounded by black curtains. Mice were allowed to individually explore the empty arena for 10 min per day of habituation. This was followed by ∼3-5 min of handling in a neighbouring room for habituation to injection procedures. TRAP2 x hM4Di-DREADD mice received additional habituation to intracerebral infusions in the form of daily handling and a dummy infusion of saline 7 days prior to activity-dependent labelling. Following habituation procedures mice remained in their experimental cages in the storage rack for at least 3 hours before being returned to their home cages with cage mates.

#### Apparatus

Behavioural testing took place in a wooden, open-topped arena (50 cm x 50 cm) with three black and one grey 40 cm high walls, positioned within a sheet-aluminium faraday cage. The floor was covered with sawdust for all experiments. Behaviour was monitored using an overhead digital camera. Objects of approximately 8 x 8 x 8 cm were constructed from Lego (Lego UK) and varied in colour and shape. The bases of the objects were secured to the arena floor using Velcro and covered with sawdust.

#### Object-in-place behavioural testing

Lego objects were placed in the four corners of the arena ∼ 10 cm from the walls. In the sample phase mice were released into the centre of the arena from their cardboard enrichment tube and allowed to explore objects for 10 min. Mice were returned to their experimental cage for a 5 min delay period during which all objects were cleaned with 100% ethanol to remove olfactory cues. Objects were then replaced into the arena with either the left- or right-hand objects exchanging position - starting positions and the side on which objects were reconfigured were counterbalanced between mice. Mice were then returned to the arena for 10 min (test phase 1) and the time spent exploring objects in novel positions was compared to exploration of objects which remained static. Either 7 days or 48h after test phase 1, mice underwent test phase 2 (10 min). Objects were placed in the arena, objects which had switched location in test phase 1 remained stationary while the objects which had been stationary in test phase 1 now switched locations (Fig. 1A). Familiar arena mice underwent the identical procedure without objects in the arena.

**Figure 1:**
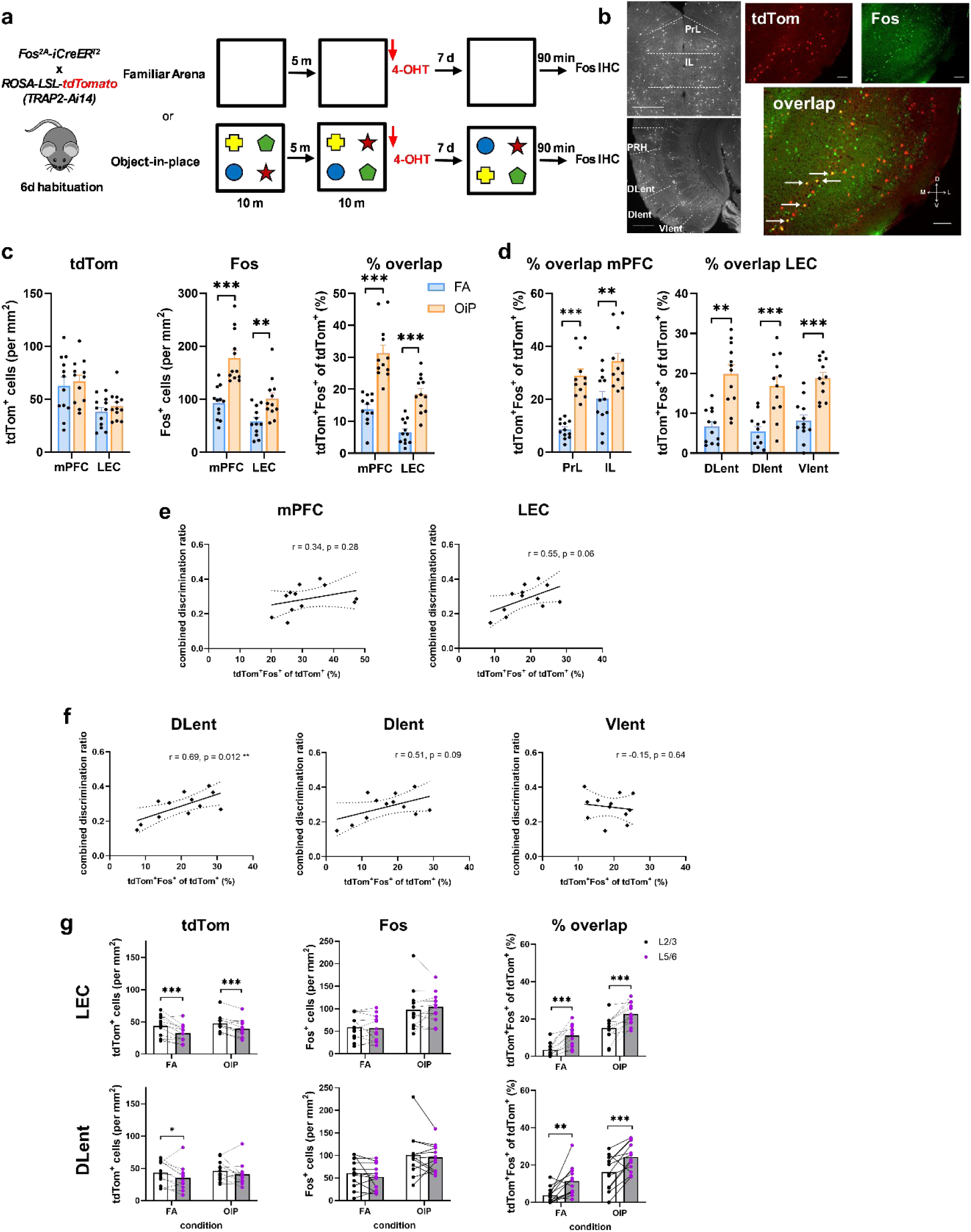
Reactivation of a dorsal lateral entorhinal cortex engram correlates with associative recognition memory performance. **a**: Experimental outline for associative memory-induced engram cell labelling. **b**: (Left) example OiP induced tdTom labelling in mPFC (upper) and LEC (lower) scale bar: 500 µm. (Right) example tdTom, Fos and overlap in LEC, arrows indicate cells expressing both tdTom and Fos; scale bar: 100 µm. **c**: (Left to right) tdTom expression, Fos expression and percent overlap in mPFC and LEC for FA and OiP behavioural conditions. **d**: Percent overlap of tdTom and Fos in mPFC subregions PrL and IL and LEC subregions DLent, DIent and VIent. **e**: Correlation between tdTom/Fos overlap and memory performance (combined discrimination ratio) in OiP behavioural task for mPFC and LEC. Dashed line show 95% confidence intervals. **f**: Same as (E) for LEC subregions. **g**: Expression levels of tdTom (left), Fos (middle) and overlap (right) in L2/3 and L5/6 of LEC (top) and DLent (bottom). Bar charts indicate mean + SEM* p < 0.05, ** p < 0.01, *** p < 0.001. N = 12 mice for all experiments.. See Supplementary Data Tables 1 & 2 for detailed statistical analysis.

#### Activity-dependent neuronal labelling

Immediately following behavioural procedures, mice were taken to the injection room, scruffed and injected intraperitoneally with 50 mg.kg^-1^ 4-hydroxytamoxifen (4-OHT; 70% Z-isomer; Hello Bio Cat#HB6040). TRAP2 x hM4Di-DREADD mice were briefly anaesthetised with isoflurane before intraperitoneal injection of 4-OHT to allow replacement of dummy cannulae (P1 technologies, Cat#C315DCS-5/SPC). 4-OHT was dissolved in 100% ethanol (10 mg.ml^-1^) at 37°C for 30 min (briefly vortexed at ∼5 min intervals), then diluted 1:2 with a 1:4 mixture of castor:sunflower seed oils (Sigma). The ethanol was evaporated off using a vacuum centrifuge and the resulting 10 mg.ml^-1^ 4-OHT solution used for injection. Mice were immediately returned to their experimental cage following injection and kept in darkness for a minimum of 3 h before being put back into home cages with cage mates and being returned to the home room. Vehicle injected animals received an injection of the castor:sunflower oil mixture at an equivalent volume to what would have been administered if they were injected with 4-OHT.

#### Chemogenetic silencing of LEC engram neurons

Following activity dependent labelling of OiP neurons, TRAP2 x hM4Di-DREADD mice were given a period of 48 h to allow expression of hM4Di protein; 48 h was chosen to attempt to restrict protein expression to neuronal somata and thus permit inhibition of only LEC engram neurons, whereas at longer delays hM4Di might be expected to accumulate in the presynaptic input to LEC neurons (Stachniak et al., 2014). Following the 48 h delay, mice received intracerebral infusions of Clozapine-N-oxide (CNO; Hello Bio Cat#HB6149) or vehicle into LEC (internal cannula, P1 technologies Cat#C315LIS-5/SPC). General infusion procedures were performed as described previously(Barker et al., 2006). CNO was dissolved at 30 µM in sterile saline and 0.05% DMSO and infused at a volume of 0.5 µl per site, over a 2 min period. Vehicle infusions consisted of sterile saline and 0.05% DMSO. 15 min after the start of infusion, mice were returned to the behavioural apparatus for 10 min for test phase 2 in which the object pair which had been static at test phase 1 exchanged position, whilst those which had been re-arranged in test phase 1 now remained static (i.e., if the left hand objects had been moved at test phase 1, the right hand objects were moved at test phase 2).

### Immunofluorescence staining

Ninety minutes after the start of test phase 2 mice were given an overdose of sodium pentobarbital (20 mg, Euthatal, Rhone Merieux) and then transcardially perfused with 0.1 M phosphate buffer (PB) followed by 4% paraformaldehyde in 0.1 M PB. Brains were removed and post-fixed in 4% paraformaldehyde for 2h and then incubated in 30% sucrose in PB at room temperature overnight. Brains were sectioned in the coronal plain into 40 µm sections with a cryostat. Four serial sets of sections were collected for each brain region of interest and stored in cryopreserve before immunofluorescent staining. Before immunostaining commenced sections were washed 5 times (2 x 30 min at RT, 3 x 10 min at 35°C) in phosphate buffered saline (PBS) to remove cryopreserve. Sections then underwent staining for Fos and tdTomato. Sections were incubated in 1% H_2_O_2_ in PBS for 30 min and then washed three times in PBS. Sections underwent antigen retrieval by incubating in sodium citrate buffer (10 mM, pH 6.0) at 80°C for 30 min. When sections had cooled down to RT they were washed four times in 0.2% Triton-X 100 in 0.1 M PBS (PBST). Sections were then incubated in mouse-on-mouse blocking buffer (Vector laboratories Cat#MKB-2213, 2 drops per 2.5 ml) in PBST for 1 h followed by incubation in blocking buffer (5% normal goat serum and 2.5% bovine serum albumin in PBST) for 1 h. Sections were then incubated in primary antibody solution to Fos (1:1000 mouse anti-Fos; Hello Bio, Cat#HB8006) and tdTomato (1:1000 chicken anti-mCherry; Abcam, Cat#ab205402) in blocking buffer for 48 h at 4°C. Sections were washed 4 times in PBST and then incubated in secondary antibody solution (1:250 goat-anti mouse Alexa Fluor 488 conjugate (Invitrogen Cat#A-11029) and 1:500 goat anti-chicken Alexa Fluor 594 conjugate (Abcam Cat#ab150176) in blocking buffer) for 24 h at 4°C. Sections were washed four times in PBST and then mounted onto polylysine coated slides. Sections were dried for 24 h in the dark and then coverslipped using Vectashield with DAPI (Vector Laboratories, Cat#H-1500).

### Electrophysiological analysis of engram TRAP2 cell excitability

48 h after activity dependent labelling of TRAP2-Ai14 mice, mice underwent test phase 2 (except Fig. 4A). Mice were then transported to a slicing room in their experimental cage covered with a black plastic bag and acute *ex vivo* brain slices were prepared. Approximately 10 minutes elapsed between the end of test 2 and the onset of slice preparation.

#### Acute slice preparation

Mice were anaesthetised with isoflurane (3%) and tested for absence of hind paw pain reflex and eye-blink reflex before decapitation. The brain was rapidly dissected and placed into a beaker of ice-cold sucrose (in mM: 189 sucrose, 26 NaHCO_3_, 10 D-glucose, 5 MgSO_4_, 3 KCl, 1.25 NaH_2_PO_4_, 0.2 CaCl_2_; Fisher Scientific) bubbled with 95% O_2_/5% CO_2_. The cerebellum was removed with a scalpel and a coronal cut was made at the approximate rostro-caudal level of the thalamus. The brain was affixed to the tissue stage of a vibratome (7000smz-2, Campden Instruments) with cyanoacrylate glue. 300 µm thick coronal brain slices were made and placed in artificial cerebrospinal fluid (aCSF; in mM: 124 NaCl, 26 NaHCO_3_, 10 D-glucose, 3 KCl, 2 CaCl_2_, 1.25 NaH_2_PO_4_, 1 MgSO_4_) bubbled with 95% O_2_/5% CO_2_. Slices were incubated for 1 hr at 34°C and then subsequently at room temperature until use. Recordings began 1 hr after slicing.

#### Electrophysiological recording

Coronal slices were placed into a submerged recording chamber perfused with 34°C aCSF at a rate of 2ml.min^-1^ and immobilised using a slice anchor (Warner Instruments). Anterior-posterior depth was determined using a brain atlas (Paxinos and Franklin, 2019) under a 4x objective and wide field oblique infra-red optics. Neurons were imaged using a 40x immersion objective and oblique infra-red and fluorescence imaging. tdTom neurons were matched with neighbouring unlabelled neurons with similar morphology in the same layer and region of the slice, typically 30-100 µm from each other. Recordings were acquired using either simultaneous dual whole-cell patch clamp or sequential single cell recordings. LEC and perirhinal cortex cellular location was determined using a brain atlas (Paxinos and Franklin, 2019); L5 neurons were 125-300 µm from the white matter, fan cells were in L2a which was visible as a dense cell layer adjacent to the almost acellular region in L1 which extended ∼100 µm from the pial surface (Vandrey et al., 2020). mPFC L2/3 and L5 were defined as 125-325 µm and 325-650 µm from the pial surface of PrL, respectively (Anastasiades et al., 2019).

Whole-cell current-clamp recordings were made using borosilicate glass micropipettes (3–6 MΩ; Harvard Apparatus, Cat# GC150-10F,) filled with potassium gluconate intracellular solution (in mM: 125 K-gluconate, 40 HEPES, 2 NaCl, 2 MgATP, 0.3 NaGTP, 0.2 EGTA, 0.25% Biocytin, 290-300 mOsm). Series resistance was 10-30 MΩ and was not compensated. Data was obtained with a Multiclamp 700A amplifier (Molecular Devices), digitized at 40 kHz (National Instruments USB-6341) and low-pass filtered at 4 kHz using WinLTP3.00 software (Anderson and Collingridge, 2007).

Resting membrane potential (RMP) was recorded immediately after break-in. Subsequently, constant-current was applied to keep cells at approximately −70 mV. Liquid junction potential was calculated to be 16.4 mV using LJPCalc and was not corrected. To assess subthreshold membrane properties and firing properties, cells were injected with a series of 500 ms square wave pulses ranging from −100 to +300 pA. To determine rheobase, cells were injected with an 800 ms linear current ramp to an amplitude sufficient to elicit at least one action potential. To evoke afterdepolarisation potentials (ADP), cells were injected with a 1-2 ms duration square current of 1000-4000 pA amplitude such that a single action potential was evoked. To evoke AHP, neurons were injected with 10 current pulses (1-2 ms duration, 1000-4000 pA amplitude) at 50 Hz. Cells were held for ∼15 min in the whole cell configuration before slow withdrawal of the recording electrode to pull an outside-out patch to aid biocytin reconstruction. The location of neurons with respect to each other was recorded to assist in identifying biocytin stained neurons.

#### Immunohistochemical biocytin staining

Following recording, slices were fixed in 4% paraformaldehyde in 0.1 M PB overnight and subsequently stored in 0.1 M PB until staining. Slices were washed 6 x 10 min in PB, then incubated in 3% H_2_O_2_ in PB for 30 min to block endogenous peroxidase activity, then washed again as above before incubation in 1% (vol/vol) avidin–biotinylated HRP complex (ABC; Vector Laboratories Cat#PK-6100) in PB containing 0.1% (vol/vol) Triton X-100 at RT for 3 hrs. Slices were then washed as above before incubation in 3,3’-diaminobenzidine solution (Sigma, Cat#D4418-50SET) for 5-10 min until staining of neuronal structures was visible. The reaction was stopped by transferring slices to cold PB, followed by further washing as above. Slices were mounted, coverslipped using Mowiol 4-88 mounting media (Sigma, Cat#81381) and allowed to dry overnight before imaging.

#### Exclusion and classification criteria

Biocytin-immunohistochemistry was used to make post-hoc corrections to cell location (layer, subregion) and morphology where neurons had been mis-labelled at the time of recording.Pyramidal neurons had primary apical dendrites projecting towards the pia mater, whilst L2 fan cells had a characteristic broad dendritic arbor extending into L1 (Tahvildari and Alonso, 2005; Vandrey et al., 2020). Neurons were excluded from analysis if their RMP exceeded −50 mV, membrane potential became persistently unstable or series resistance exceeded 30 MΩ. Individual electrophysiological sweeps were excluded where membrane potential was transiently unstable, deviated from −70 mV, or spontaneous activity prevented accurate measurement of data.

### Quantification and statistical analysis

#### Behaviour scoring and statistics

Exploratory behaviour was manually scored and defined as the animal directing its nose towards the object at a distance <2 cm and actively exploring the object. Any other behaviour, such as looking around while sitting on or resting against the object, grooming within 2 cm of an object, or digging at object bases, was not considered object exploration. Discrimination between the objects was calculated using a discrimination ratio (DR), calculated as the absolute difference in the time spent exploring the moved and unmoved stimuli divided by the time spent exploring all the objects. The DR takes into account individual differences in the total amount of object exploration (Ennaceur and Delacour, 1988; Dix and Aggleton, 1999). For the combined DR the exploration of the moved stimuli in test phase one and two was added together and the same was done for the unmoved stimuli and these combined values were used to calculate the combined DR. Performance in each test phase and for the combined DR was assessed using a two-tailed one-sample t-test against chance (DR = 0). Exploration across the phases of the task were compared using a one-way within subjects ANOVA. Post-hoc comparisons used a Bonferroni correction. For chemogenetic silencing of engram cells in LEC, DR and total object exploration levels across test phase 1 and 2 was compared using a two-way ANOVA with drug as a between subjects factor and test phase as a within subjects factor. Post-hoc comparisons examined simple main effects. Comparison of exploration in the sample phase used a one-way between subjects ANOVA. Performance in the test phases was assessed against chance using a one-sample t-test as previously described.

#### Cell counting protocol and statistics

For the quantification of tdTom, Fos and DAPI expression in mPFC, LEC and PRH, cells were counted bilaterally from 4 (mPFC) or 5 (LEC and PRH) coronal sections from specific locations along the anterior-posterior length of each brain region (Table 1). Fluorescent images were acquired using a Leica Microsystems DM6 B microscope and Hamamatsu C13400-20CU Digital CMOS camera via the Leica Application Suite X software using 100x magnification. Manual cell counting was performed using ImageJ software. Images were split into individual channels. For each cortical area, a region of interest (ROI; 0.5 mm height by 0.25 mm width) was placed over layer 2/3 and layer 5/6 using the DAPI channel in each slice. Cortical regions were identified using a brain atlas (Paxinos and Franklin, 2019). Cell counts for each channel were obtained as follows; background subtraction was applied to remove background noise, thresholds were set and images were converted to binary. The analyse particles tool was used to quantify the number of labelled cells in each ROI based on particle size (50-500 µm^2^ tdTom, 20-150 µm^2^ Fos) and circularity (0.4-1.0). Overlap between tdTom and Fos expression was determined by digitally merging the analysed images for each ROI to form a composite image and the number of co-labelled cells quantified. All cell counting was performed by an experimenter blind to the behavioural condition of each animal.

**Table 1:** Anterior-posterior position relative to bregma of coronal sections analysed for tdTom and Fos expression and overlap. PrL prelimbic cortex, IL infralimbic cortex, PRH perirhinal cortex, DLent dorsal lateral entorhinal cortex, DIent dorsal intermediate entorhinal cortex, VIent ventral intermediate entorhinal cortex.

| Section | AP position, relative to bregma | Brain regions analysed |
| --- | --- | --- |
| mPFC 1 | +2.33mm to +1.97mm | PrL |
| mPFC 2 | +1.97mm to +1.77mm | PrL, IL |
| mPFC 3 | +1.77mm to +1.41mm | PrL, IL |
| mPFC 4 | +1.41mm to +1.33mm | IL |
| LEC 1 | -2.91mm to -3.15mm | PRH, DLent, Dlent |
| LEC 2 | -3.15mm to -3.27mm | PRH, DLent, Dlent |
| LEC 3 | -3.39mm to -3.63mm | PRH, DLent, Dlent, Vlent |
| LEC 4 | -3.79mm to -4.10mm | PRH, DLent, Dlent, Vlent |
| LEC 5 | -4.23mm to -4.47mm | PRH, DLent, Dlent, Vlent |

To analyse each brain subregion, cell counts in L2/3 and L5/6 ROIs were combined for each hemisphere, the counts for each hemisphere and for each AP depth counted were averaged. To generate counts for all of mPFC and LEC, the counts for the individual subregions were averaged in each animal to generate counts for the whole brain region. To generate counts for L2/3 and L5/6 of cortex, counts in each region were averaged across hemispheres and sections. These averaged counts were then converted into counts per mm^2^. Percent overlap was calculated as (tdTom^+^Fos^+^ cells/tdTom^+^ cells) * 100.

Comparisons of cell counts across the whole cortical regions between behavioural conditions used a one-way ANOVA with behavioural condition as a between subject factor. Comparison of layer 2/3 and layer 5/6 counts between behavioural conditions used a two-way ANOVA with cortical depth as a within-subjects factor and behavioural condition as a between-subjects factor. Post-hoc tests analysed simple main effects where appropriate. To analyse the relationship between memory performance and cell reactivation a Pearson’s correlation coefficient was calculated comparing combined DR and % reactivation. All statistical analyses were performed using SPSS and significance was set at p < 0.05.

#### Electrophysiology statistics

Electrophysiological data were imported into MATLAB for analysis using custom-written scripts based upon Kerrigan et al. 2014 (code available on request). Action potential parameters are measured from the first action potential from the smallest square-wave current-injection which yielded an action potential. Reported input resistance values are steady-state input resistance. Statistical analysis of all electrophysiological variables was performed in GraphPad Prism 10 using Mann Whitney U test, except action potential firing where a mixed-effect model with a Greenhouse-Geisser correction applied and labelling and current injection as factors was used, and ADP presence where a Chi-squared test was used. Cells were considered the experimental unit for electrophysiology.

## Results

### OiP memory performance correlates with reactivation of dorsolateral entorhinal cortex engram cells

TRAP2 mice were crossed with tdTomato (tdTom) Cre reporter *Ai14* mice (Madisen et al., 2010) resulting in Fos-driven, 4-hydroxytamoxifen (4-OHT)-dependent tdTom labelling. Whilst rewarding and aversive experiences produce high levels of labelling and reactivation at recall in TRAP mice (DeNardo et al., 2019; Bobadilla et al., 2020), it is unknown whether every day, spontaneous learning behaviours such as associative recognition memory are processed in the same manner. TRAP2-Ai14 littermates were assigned to one of two experimental groups: OiP or familiar arena (FA), the latter of which served as the control group (Fig. 1A). Mice assigned to the OiP group preferentially explored reconfigured objects when tested at 5 min and 7 d, demonstrating intact memory (Fig. S1A).

We focussed our analysis of TRAP2 cell activation and reactivation on two cortical regions known to be essential for associative recognition memory: LEC (Wilson et al., 2013; Vandrey et al., 2020; Tozzi et al., 2024; Kinnavane et al., 2025) and mPFC (Preston and Eichenbaum, 2013; Barker et al., 2017). Mice which had not received 4-OHT showed negligible tdTom labelling (Fig. S1B). Analysis of cell activation patterns across the whole of mPFC and LEC (Fig. 1B) revealed the number of tdTom labelled cells was not significantly different between the OiP and FA conditions in either brain region (Fig. 1B, Table S1). However, the number of Fos immunostained cells following recall at 7 d was significantly higher in OiP compared to FA mice and, critically, the percentage of Fos-immunostained tdTom cells (indicating reactivation) was significantly higher in the OiP condition compared to FA in both brain regions (Fig. 1C, Table S1). These same patterns of activation and reactivation were also seen when mPFC was separated into prelimbic (PrL) and infralimbic cortex (IL) subdivisions and the LEC into dorsal lateral (DLent), dorsal intermediate (DIent) and ventral intermediate (VIent) subdivisions (Fig. 1D; S1C, Table S1).

To investigate the relationship between memory performance in the OiP task and engram cell reactivation, a Pearson correlation coefficient was computed for each brain region. No significant correlation was found between the combined discrimination ratio (DR) and tdTom cell reactivation in mPFC, nor the PrL and IL subdivisions (Fig. 1E, S1D). Analysis of LEC showed a non-significant trend towards positive correlation between memory performance and tdTom cell reactivation (Fig. 1E). Interestingly, analysis of LEC subdivisions identified a statistically significant positive correlation in DLent, a positive but not statistically significant correlation in DIent and a non-significant negative relationship in VIent (Fig. 1E). DLent may, therefore, be a critical region for OiP memory retrieval.

The superficial and deep layers of LEC are anatomically distinct, with layers 2 & 3 (L2/3) providing input into the hippocampus while layers 5 & 6 (L5/6) receive inputs from the hippocampus and project to non-hippocampal regions (Ohara et al., 2021). In the whole LEC significantly more tdTom cells were found in L2/3 than L5/6 in both the FA and OiP conditions, whereas memory retrieval produced equal amounts of Fos labelling in L2/3 and L5/6 in both behavioural tasks (Fig. 1G). Surprisingly, a significantly higher proportion of deep layer LEC tdTom cells were reactivated by retrieval in both behavioural conditions (Fig. 1G). Analysis of LEC subdivisions showed that this activation and reactivation pattern was consistent across all three LEC subdivisions (Fig. 1G, S2A,B, Table S2), with the exception of there being no difference in tdTom labelling in different layers of DLent of OiP mice (Fig. 1G, Table S2) or in the reactivation of VIent layers in OiP mice (Fig. S2B, Table S2).

Layer analysis of mPFC as a whole did not reveal any differences in labelling between L2/3 and L5/6 (Fig. S2C). In PrL, significantly more tdTom cells were found in L2/3 in both behavioural tasks, however there was no significant difference in Fos labelling or reactivation between layers following memory retrieval (Fig. S2D). In IL significantly more tdTom cells were activated in L5/6 in the OiP task, however no significant differences in layer were observed for Fos labelling or reactivation following memory retrieval for either behavioural condition (Fig. S2E, Table S2).

In summary these data show that OiP recall robustly reactivates LEC and mPFC neurons which were activated at encoding, that reactivation of LEC neurons correlates with memory performance with a dorsoventral gradient, and that reactivation is highest in L5/6 LEC neurons, putatively identifying deep layer DLent as a key locus of object-place representations. We next determined the necessity of LEC TRAP2 neurons for memory performance.

### Inactivation of LEC TRAP2 cells impairs OiP memory recall

If activity-labelled neurons are to be considered engram cells, in addition to their reactivation by stimuli relevant to the labelling experience, their activity should also be necessary for retrieval of the memory (Tanaka et al., 2014). To test this hypothesis, TRAP2 mice were crossed with Cre-dependent *hM4Di-DREADD* mice to permit chemogenetic inactivation of activated neurons (Fig. 2A). Guide cannulae were implanted bilaterally in LEC (Fig. 2B), targeting DLent where TRAP2 cell reactivation was significantly correlated with memory performance (Fig. 1F). Following recovery from surgery, mice underwent OiP testing before injection with 4-OHT to drive expression of hM4Di in active neurons (Fig. 2A). We then assessed the effect of hM4Di-mediated inactivation on memory retrieval by infusing vehicle or CNO into LEC prior to the second test phase, 48 h after labelling neurons.

**Figure 2:**
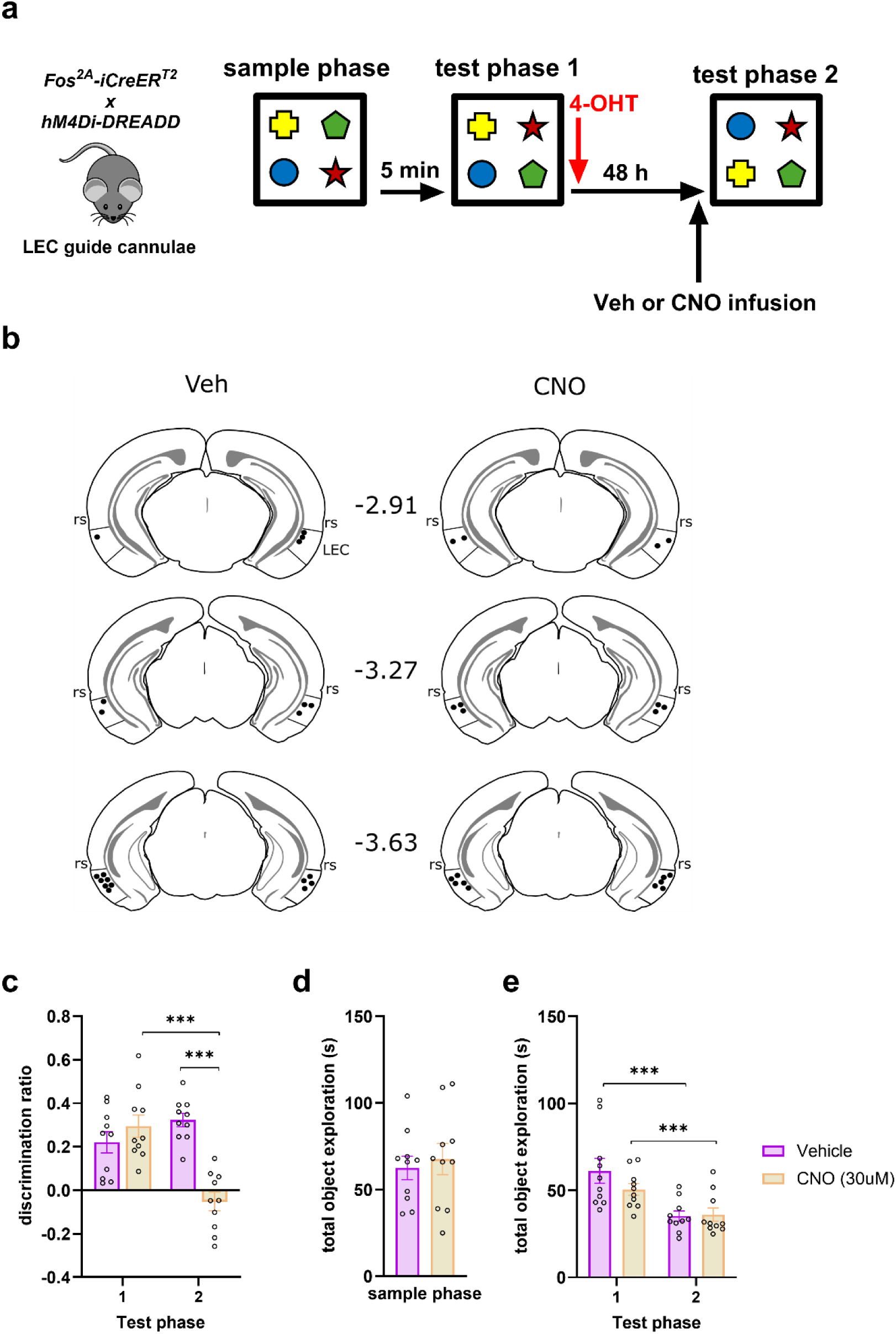
Engram cells in lateral entorhinal cortex are essential for the retrieval of associative recognition memory. **a**: Experimental outline for deactivating LEC engram cells during associative recognition memory retrieval. **b**: LEC cannula locations (represented by black dots) in the vehicle (left) and CNO (right) infused animals. Numbers represent AP position in mm relative to bregma. rs = rhinal sulcus. **c**: Memory performance (discrimination ratio) at test phase 1 and test phase 2 of vehicle and CNO infused animals. **d**: Total object exploration during the sample phase of the OiP task. **e**: Total object exploration levels during test phase 1 and test phase 2 of the object in place task. Bar charts show mean ± SEM. *** p<0.001. N = 10 mice for all conditions. For statistical analysis see text.

Mice assigned to both vehicle and CNO groups showed intact memory at the 5 min delay (Fig. 2C, vehicle t_(9)_ = 4.53, p = 0.001; CNO t_(9)_ = 5.73, p = 0.0003). Critically, however, mice infused with CNO prior to testing at 48 h had impaired memory, while vehicle infused mice had intact memory (Fig. 2C, Drug by test phase F_(1,18)_ = 21.92, p = 0.0002; vehicle: t_(9)_ = 10.39, p = 3.0 x 10^-6^; CNO: t_(9)_ = −1.24, p = 0.12). No difference in exploration levels were found between vehicle and CNO groups at the sample (Fig. 2D, F_(1,18)_ = 0.21 p = 0.656) or either test phase (Fig. 2E, drug by test phase F_(1,18)_ = 4.50, p = 0.048; vehicle vs CNO, test 1: p = 0.194, test 2: p = 0.862; T1 vs T2, vehicle: p = 2×10^-6^, CNO: p = 0.001). These data demonstrate the necessity of LEC TRAP neurons for successful retrieval of OiP memory, therefore definitively demonstrating their identity as engram neurons.

### Object reconfiguration increases excitability of L5 LEC TRAP2 neurons

Material changes in engram neurons are hypothesised to increase the likelihood of their reactivation upon exposure to specific cues. One mechanism by which this can be achieved is by changes in cellular excitability, which may facilitate firing in response to complete, partial or ambiguous cues (Cai et al., 2016; Whitaker et al., 2017a; Pignatelli et al., 2019) and support consolidation (Thompson et al., 1996; Zhang and Linden, 2003).

We investigated neuronal excitability in *TRAP2-Ai14* mice 48 h after TRAP labelling by preparing *ex vivo* slices immediately after memory recall (Fig. 3A), focussing on deep-layer LEC neurons, which had shown the highest levels of reactivation by OiP memory recall (Fig. 1G). In slices from these mice, tdTom L5 pyramidal neurons exhibited increased firing in response to depolarising current compared to unlabelled neurons (Fig. 3B). Further analysis of the physiological properties of these neuronal populations (Fig. 3B; Table 2) showed input resistance was significantly higher in tdTom neurons. Rheobase, a measure related to input resistance and action potential threshold, was lower in tdTom neurons but not statistically different, likely owing to an insignificant trend to a more depolarised action potential threshold in tdTom neurons. Other membrane properties that influence firing, including afterhyperpolarisation (AHP) potentials (Disterhoft and Oh, 2006; Whitaker et al., 2017b), were not significantly different (Fig. 3B; Table 2). In addition to increased excitability, tdTom neurons also showed modified action potential dynamics, exhibiting lower amplitude action potentials which had a slower rate of rise than those observed in unlabelled neurons (Fig. 3B; Table 2). Collectively these data show that exposure to a novel configuration of engram-related objects elicits changes in the electrophysiological properties of tdTom neurons, resulting in increased excitability and firing rate in response to depolarising stimuli.

**Figure 3.**
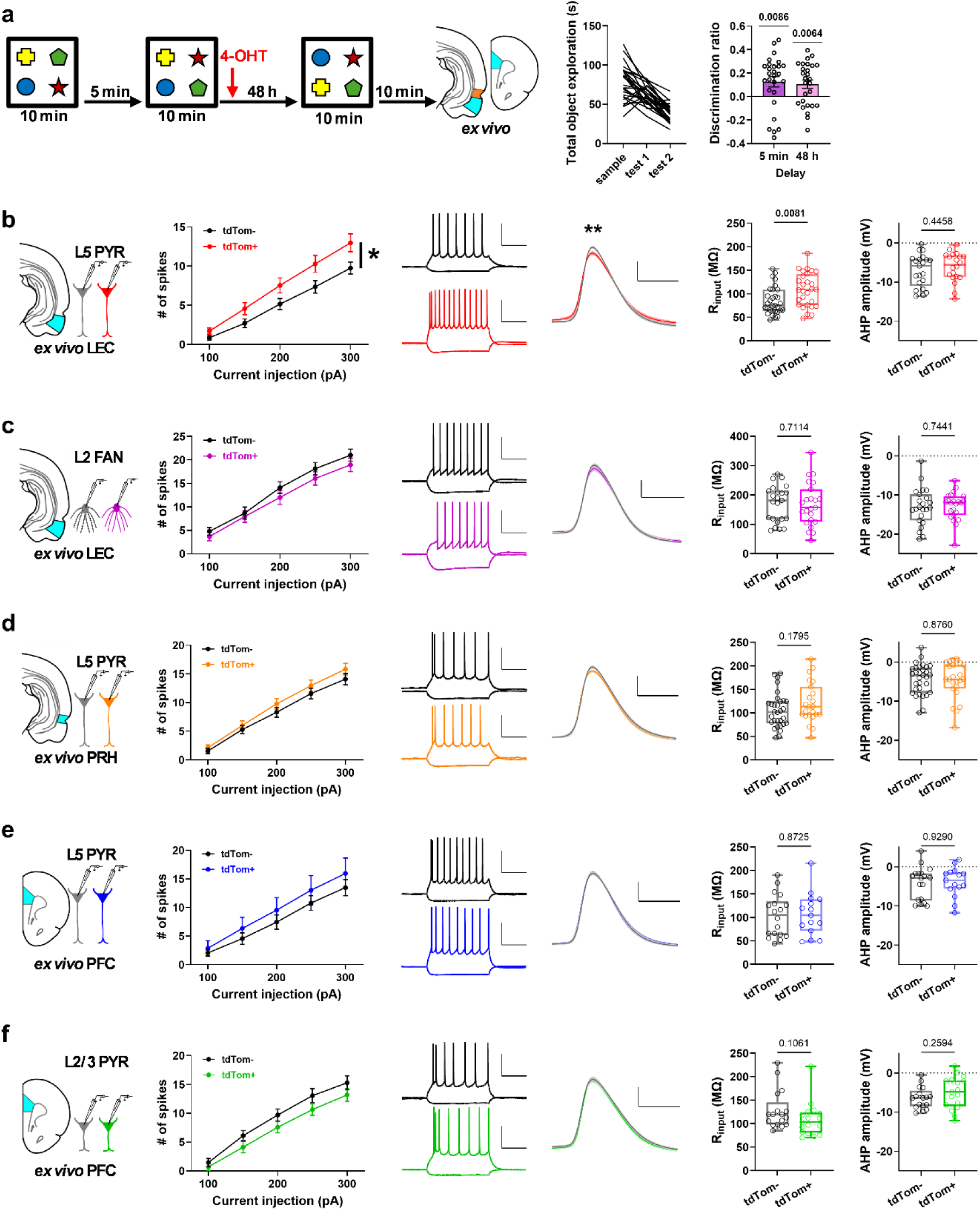
Object reconfiguration increases excitability of LEC L5 pyramidal engram cells, but not other cortical engram cells. **a**: (Left) Experimental schematic for *ex vivo* recordings from animals exposed to reconfigured objects before slicing. (Right) exploration and discrimination ratio, bars represent mean ± SEM (n = 27 mice). **b**: Left to right: action potential firing (mean ± SEM), representative voltage responses to −100 and +200 pA current injections for a tdTom^-^ and tdTom^+^ cell pair, average action potential waveforms, input resistance and afterhyperpolarisation (AHP) amplitude of LEC L5 pyramidal neurons. Boxplots show median (centre line), 25^th^ and 75^th^ percentiles, whiskers show minimum and maximum points. tdTom^-^ n = 31 cells from 16 mice, tdTom^+^ n = 28 cells from 16 mice. **c**: Same as (**b**) but for L2 fan cells. tdTom^-^ n = 23 cells from 16 mice, tdTom^+^ n = 23 cells from 17 mice. **d**: Same as (**b**) but for perirhinal cortex L5 pyramidal cells. tdTom^-^ n = 31 cells from 14 mice, tdTom^+^ n = 23 cells from 13 mice. **e**: Same as (**b**) but for prefrontal cortex L5 pyramidal cells. tdTom^-^ n = 18 cells from 5 mice, tdTom^+^ n = 15 cells from 5 mice. **f**: Same as (**b**) but for prefrontal cortex L2/3 pyramidal cells. tdTom^-^ n = 17 cells from 5 mice, tdTom^+^ n = 17 cells from 5 mice. Scale bars 200 ms, 40 mV. * = p<0.05, ** = p < 0.01. See Supplementary Data Tables 3-5 for further parameters and statistical analysis.

**Table 2.**
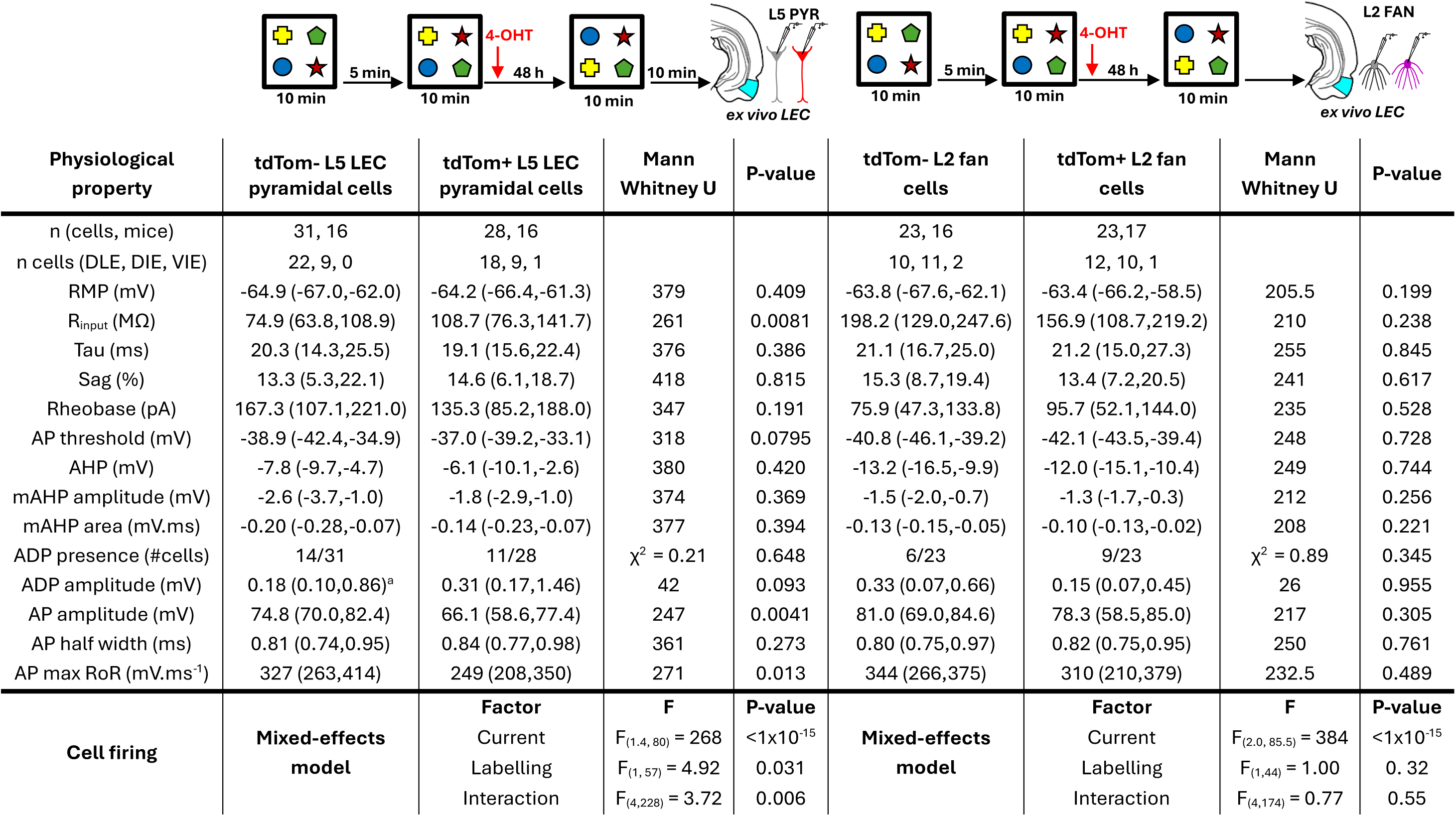
Summary of physiological properties of LEC layer 5 pyramidal and layer 2 fan tdTom- and tdTom+ neurons recorded from ex vivo slices from animals which underwent a second test phase (related to Fig. 3b,c). Values represent median (25,75 percentiles). Reported statistical values show Mann Whitney U test with 2-tailed p-value, except for ADP presence which uses Chi-squared test with 2-tailed p value. ^a^*one cell was excluded from analysis due to spiking during ADP*. RMP resting membrane potential, R_input_ input resistance, AP action potential, AHP afterspike-hyperpolarisation, mAHP medium AHP, ADP afterspike-depolarisation, RoR rate of rise.

| Physiological property | tdTom- L5 LEC pyramidal cells | tdTom+ L5 LEC pyramidal cells | Mann Whitney U | P-value | tdTom- L2 fan cells | tdTom+ L2 fan cells | Mann Whitney U | P-value |
| --- | --- | --- | --- | --- | --- | --- | --- | --- |
| n (cells, mice) | 31, 16 | 28, 16 |  |  | 23, 16 | 23,17 |  |  |
| n cells (DLE, DIE, VIE) | 22, 9, 0 | 18, 9, 1 |  |  | 10, 11, 2 | 12, 10, 1 |  |  |
| RMP (mV) | -64.9 (-67.0,-62.0) | -64.2 (-66.4,-61.3) | 379 | 0.409 | -63.8 (-67.6,-62.1) | -63.4 (-66.2,-58.5) | 205.5 | 0.199 |
| R<sub>input</sub> (MΩ) | 74.9 (63.8,108.9) | 108.7 (76.3,141.7) | 261 | 0.0081 | 198.2 (129.0,247.6) | 156.9 (108.7,219.2) | 210 | 0.238 |
| Tau (ms) | 20.3 (14.3,25.5) | 19.1 (15.6,22.4) | 376 | 0.386 | 21.1 (16.7,25.0) | 21.2 (15.0,27.3) | 255 | 0.845 |
| Sag (%) | 13.3 (5.3,22.1) | 14.6 (6.1,18.7) | 418 | 0.815 | 15.3 (8.7,19.4) | 13.4 (7.2,20.5) | 241 | 0.617 |
| Rheobase (pA) | 167.3 (107.1,221.0) | 135.3 (85.2,188.0) | 347 | 0.191 | 75.9 (47.3,133.8) | 95.7 (52.1,144.0) | 235 | 0.528 |
| AP threshold (mV) | -38.9 (-42.4,-34.9) | -37.0 (-39.2,-33.1) | 318 | 0.0795 | -40.8 (-46.1,-39.2) | -42.1 (-43.5,-39.4) | 248 | 0.728 |
| AHP (mV) | -7.8 (-9.7,-4.7) | -6.1 (-10.1,-2.6) | 380 | 0.420 | -13.2 (-16.5,-9.9) | -12.0 (-15.1,-10.4) | 249 | 0.744 |
| mAHP amplitude (mV) | -2.6 (-3.7,-1.0) | -1.8 (-2.9,-1.0) | 374 | 0.369 | -1.5 (-2.0,-0.7) | -1.3 (-1.7,-0.3) | 212 | 0.256 |
| mAHP area (mV.ms) | -0.20 (-0.28,-0.07) | -0.14 (-0.23,-0.07) | 377 | 0.394 | -0.13 (-0.15,-0.05) | -0.10 (-0.13,-0.02) | 208 | 0.221 |
| ADP presence (#cells) | 14/31 | 11/28 | $\chi^2 = 0.21$ | 0.648 | 6/23 | 9/23 | $\chi^2 = 0.89$ | 0.345 |
| ADP amplitude (mV) | 0.18 (0.10,0.86)<sup>a</sup> | 0.31 (0.17,1.46) | 42 | 0.093 | 0.33 (0.07,0.66) | 0.15 (0.07,0.45) | 26 | 0.955 |
| AP amplitude (mV) | 74.8 (70.0,82.4) | 66.1 (58.6,77.4) | 247 | 0.0041 | 81.0 (69.0,84.6) | 78.3 (58.5,85.0) | 217 | 0.305 |
| AP half width (ms) | 0.81 (0.74,0.95) | 0.84 (0.77,0.98) | 361 | 0.273 | 0.80 (0.75,0.97) | 0.82 (0.75,0.95) | 250 | 0.761 |
| AP max RoR (mV.ms<sup>-1</sup>) | 327 (263,414) | 249 (208,350) | 271 | 0.013 | 344 (266,375) | 310 (210,379) | 232.5 | 0.489 |
| **Cell firing** | **Mixed-effects model** | **Factor** | **F** | **P-value** | **Mixed-effects model** | **Factor** | **F** | **P-value** |
| | | Current | $F_{(1.4, 80)} = 268$ | $<1 \times 10^{-15}$ | | Current | $F_{(2.0, 85.5)} = 384$ | $<1 \times 10^{-15}$ |
| | | Labelling | $F_{(1, 57)} = 4.92$ | 0.031 | | Labelling | $F_{(1, 44)} = 1.00$ | 0.32 |
| | | Interaction | $F_{(4, 228)} = 3.72$ | 0.006 | | Interaction | $F_{(4, 174)} = 0.77$ | 0.55 |

### Excitability changes were not observed in other cortical engram cells

That tdTom L5 LEC pyramidal neurons are both reactivated and show heightened excitability following exposure to rearranged objects suggests a key role in OiP memory. We next asked whether other neuronal populations activated by OiP memory also undergo changes in cellular excitability. tdTom labelling was found to be abundant in LEC L2a fan cells, which are required for episodic-like (Vandrey et al., 2020) and associative memory (Lee et al., 2021), in *ex vivo* slices from mice exposed to reconfigured objects. However, unlike L5 pyramidal neurons, no differences in spiking or other measures of intrinsic excitability or action potential waveforms were found between tdTom and unlabelled L2a fan cells (Fig. 3C, Table 2).

We next examined excitability of pyramidal neurons in PRH, a major input to LEC which encodes object identity and novelty (Seoane et al., 2012) but which, in contrast to LEC, is not thought to encode object-location associations (Deshmukh et al., 2012). Labelling in PRH had a similar pattern to that seen in in LEC: both Fos labelling (Fig. S1C) and tdTom cell reactivation (Fig. 1E) after retrieval were elevated in OiP mice compared to FA, although reactivation of PRH engram cells was not significantly correlated with memory performance (Fig. S1F). tdTom and Fos labelling was higher in PRH L5/6 than L2/3 (Fig. S2F) irrespective of task, and reactivation was enhanced in L5/6 compared to L2/3 in OiP mice only. However, no differences in action potential firing or other electrophysiological parameters between tdTom and unlabelled PRH neurons were observed (Fig. 3D, Table 3).

**Table 3.**
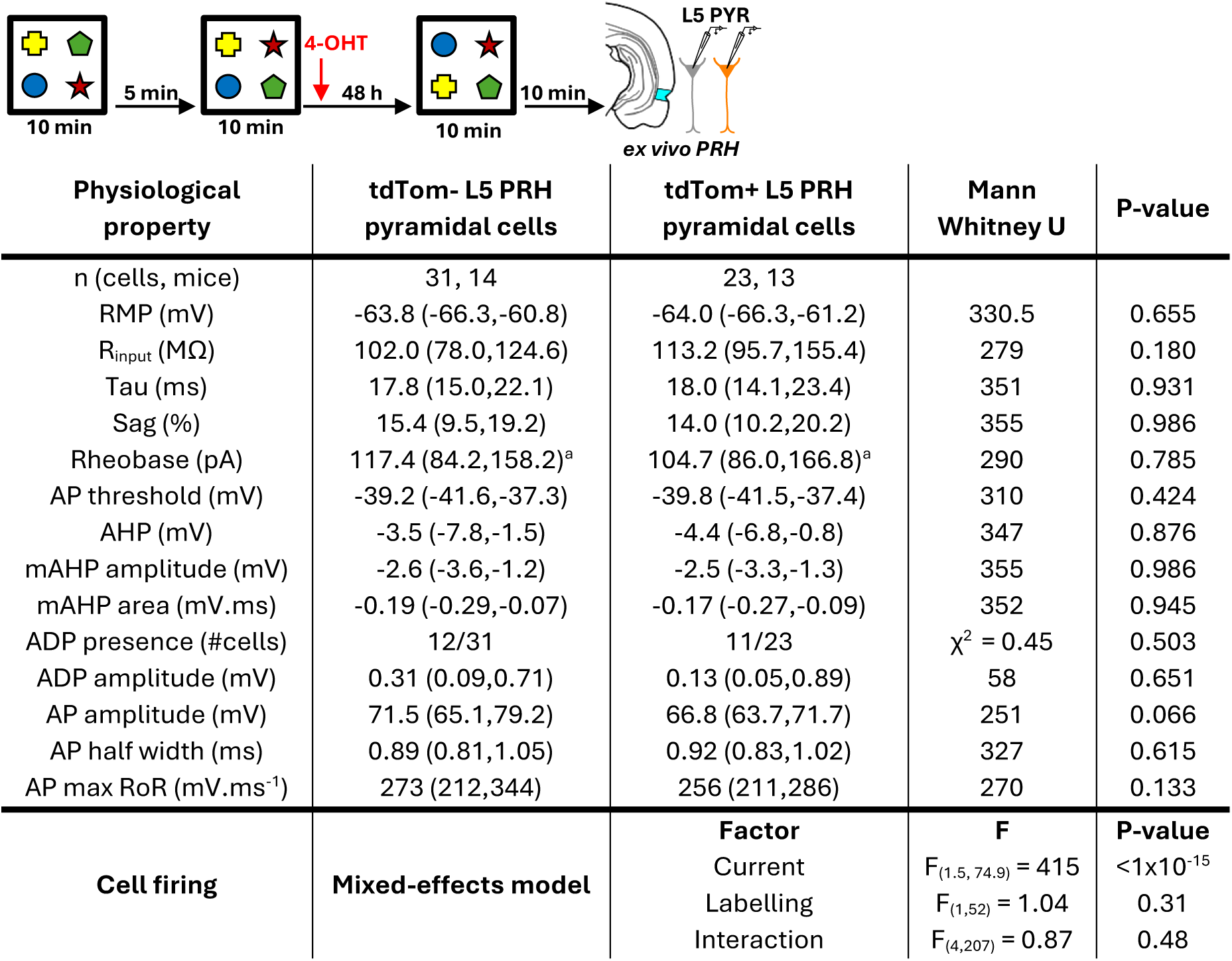
Summary of physiological properties of layer 5 perirhinal cortex tdTom- and tdTom+ neurons recorded from ex vivo slices from animals which underwent a second test phase (related to Fig. 3d). Values represent median (25,75 percentiles). Reported statistical values show Mann Whitney U test with 2-tailed p-value, except for ADP presence which uses Chi-squared test with 2-tailed p value. ^a^*data not collected for 2 cells*. RMP resting membrane potential, Rinput input resistance, AP action potential, AHP afterspike-hyperpolarisation, mAHP medium AHP, ADP afterspike-depolarisation, RoR rate of rise

We additionally assessed excitability in mPFC, however no changes in the firing rate nor other electrophysiological measures were found in L5 pyramidal cells (Fig. 3E, Table 4) or L2/3 pyramidal cells (Fig. 3F, Table 4). Together these data show that whilst OiP recall results in significant reactivation of tdTom labelled neurons in multiple cortical regions and layers, increases in cellular excitability and action potential dynamics were only found in L5 LEC neurons.

**Table 4.** Summary of physiological properties of layer 5 and 2/3 prefrontal cortex tdTom- and tdTom+ pyramidal neurons recorded from ex vivo slices from animals which underwent a second test phase (related to Fig. 3e,f). Values represent median (25,75 percentiles). Reported statistical values show Mann Whitney U test with 2-tailed p-value, except for ADP presence which uses Chi-squared test with 2-tailed p value. ^a^*data not collected for 1 cell*. ^b^*data not collected for 3 cells*. ^c^*data not collected for 4 cells*. RMP resting membrane potential, R_input_ input resistance, AP action potential, AHP afterspike-hyperpolarisation, mAHP medium AHP, ADP afterspike-depolarisation, RoR rate of rise

| Physiological property | tdTom- L5 mPFC pyramidal cells | tdTom+ L5 mPFC pyramidal cells | Mann Whitney U | P-value | tdTom- L2/3 PFC pyramidal cells | tdTom+ L2/3 PFC pyramidal cells | Mann Whitney U | P-value |
| --- | --- | --- | --- | --- | --- | --- | --- | --- |
| n (cells, mice) | 18, 5 | 18, 5 |  |  | 17, 5 | 17, 5 |  |  |
| RMP (mV) | -64.2 (-66.2,-63.1) | -64.9 (-67.4,-61.8) | 127 | 0.782 | -64.3 (-66.1,-62.3) | -63.4 (-66.2,-60.9) | 133 | 0.702 |
| R<sub>input</sub> (MΩ) | 105.5 (62.8,133.9) | 104.6 (72.0,138.8) | 130 | 0.873 | 119.7 (98.1,145.4) | 103.4 (80.5,123.5) | 97 | 0.106 |
| Tau (ms) | 13.4 (9.4,18.4) | 13.8 (9.1,15.2) | 133 | 0.957 | 16.0 (12.8,17.9) | 15.0 (12.4,19.3) | 139 | 0.865 |
| Sag (%) | 7.9 (5.1,11.3) | 4.4 (2.3,13.3) | 110 | 0.381 | 3.5 (-2.7,7.6) | 2.3 (-2.6,4.1) | 122 | 0.454 |
| Rheobase (pA) | 109.5 (64.4,143.5)<sup>a</sup> | 117.7 (85.1,171.9)<sup>a</sup> | 99 | 0.444 | 99.4 (82.7,130.0) | 137.6 (99.4,172.7)<sup>a</sup> | 98 | 0.179 |
| AP threshold (mV) | -38.7 (-44.1,-37.7) | -39.6 (-41.2,-38.3) | 133 | 0.957 | -36.6 (-39.0,-34.8) | -37.9 (-40.4,-35.4) | 114 | 0.306 |
| AHP (mV) | -2.8 (-8.6,-1.7) | -3.5 (-5.6,-1.4) | 132 | 0.929 | -6.4 (-8.5,-4.5) | -4.8 (-8.5,-1.9) | 111 | 0.260 |
| mAHP amplitude (mV) | -2.1 (-2.7,-1.1)<sup>c</sup> | -1.1 (-3.1,-0.5)<sup>b</sup> | 83 | 0.720 | -0.8 (-2.0,-0.4) | -1.1 (-2.3,-0.3)<sup>a</sup> | 131 | 0.873 |
| mAHP area (mV.ms) | -0.15 (-0.18,-0.07)<sup>c</sup> | -0.08 (-0.19,-0.03)<sup>b</sup> | 80 | 0.616 | -0.06 (-0.15,-0.02) | -0.09 (-0.19,-0.02)<sup>a</sup> | 131 | 0.873 |
| ADP presence (#cells) | 8/18 | 9/15<sup>b</sup> | $\chi^2 = 0.79$ | 0.373 | 10/17 | 7/17 | $\chi^2 = 1.06$ | 0.303 |
| ADP amplitude (mV) | 0.94 (0.32,1.70) | 0.79 (0.12,1.56) | 40 | 0.677 | 0.32 (1.13,1.03) | 0.22 (0.04,1.02) | 19 | 0.535 |
| AP amplitude (mV) | 67.9 (64.0,79.5) | 68.9 (65.5,76.2) | 134 | 0.986 | 65.2 (58.0,78.9) | 69.6 (62.7,76.8) | 131 | 0.658 |
| AP half width (ms) | 0.88 (0.80,0.98) | 0.92 (0.75,1.01) | 123 | 0.682 | 0.99 (0.91,1.07) | 0.93 (0.85,1.00) | 114 | 0.306 |
| AP max RoR (mV.ms<sup>-1</sup>) | 271 (244,349) | 266 (236,323) | 131 | 0.901 | 238 (209,338) | 269 (235,344) | 117 | 0.357 |
| **Cell firing** | **Mixed-effects model** | **Factor** | **F** | **P-value** | **Mixed-effects model** | **Factor** | **F** | **P-value** |
| | | Current | $F_{(1.2, 37.7)} = 144$ | $<1 \times 10^{-15}$ | | Current | $F_{(1.5, 46.4)} = 424$ | $<1 \times 10^{-15}$ |
| | | Labelling | $F_{(1, 31)} = 0.68$ | 0.41 | | Labelling | $F_{(1, 32)} = 2.04$ | 0.16 |
| | | Interaction | $F_{(4, 123)} = 0.58$ | 0.68 | | Interaction | $F_{(4, 127)} = 1.6$ | 0.18 |

### Increased neuronal excitability of TRAP2 LEC L5 pyramidal neurons requires recall of engram-specific cues with associated novel information

We next examined the behavioural conditions required to elicit changes in engram cell excitability. Increased cellular excitability has been suggested as a mechanism either leading to, or resulting from, recruitment to an engram (Disterhoft and Oh, 2006; Josselyn and Frankland, 2018); if such changes persist to 48 h after encoding, then these may explain the increased excitability of LEC L5 tdTom neurons. We investigated this possibility by recording from LEC L5 pyramidal neurons 48 h after encoding, without recall prior to slicing (Fig. 4A). We found that tdTom and unlabelled cells had equal rates of firing, cellular excitability and action potential properties (Fig. 4A, Table 5). These data show that increased excitability of tdTom LEC L5 pyramidal cells requires re-exposure to the engram objects.

**Figure 4.**
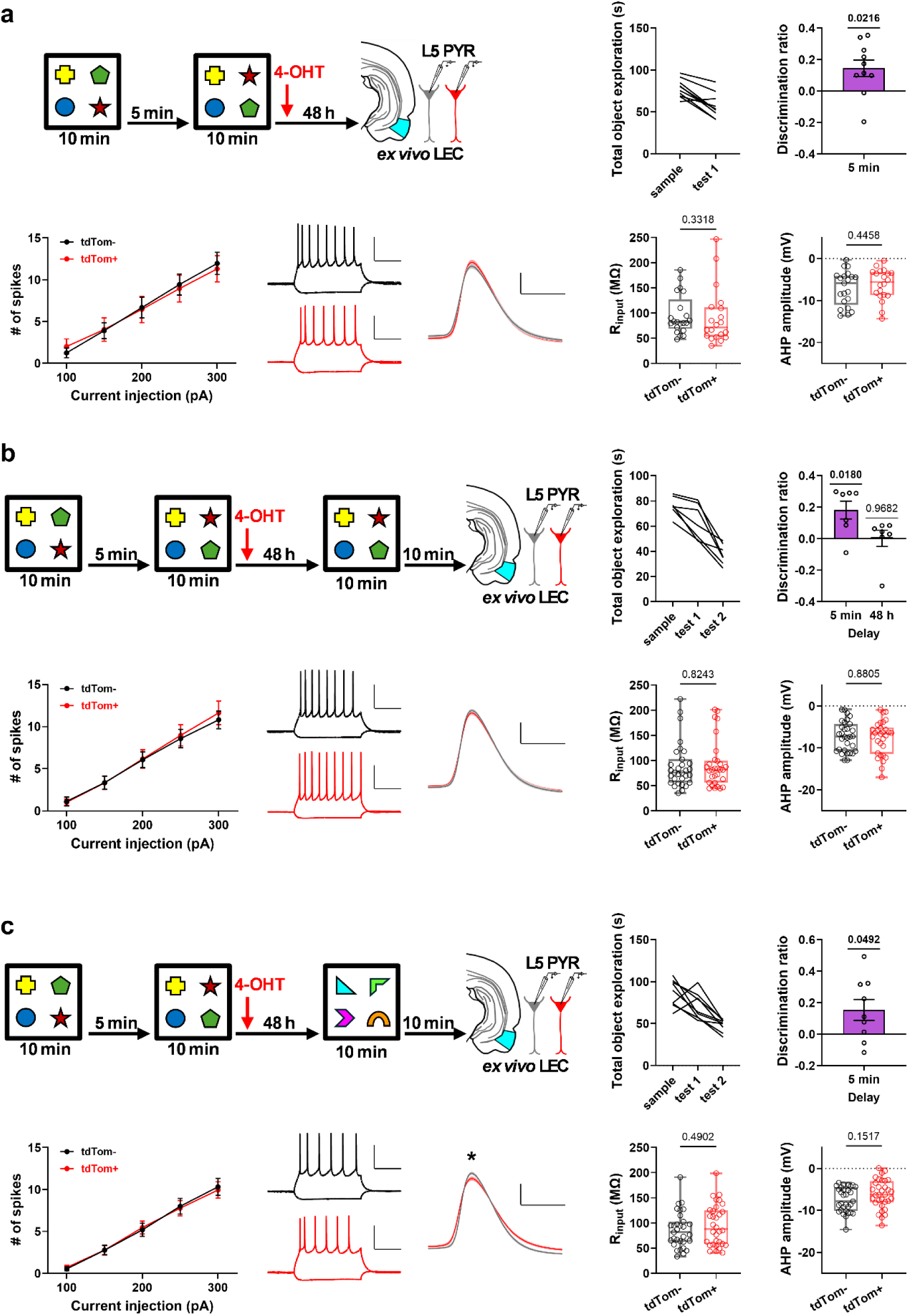
Increased neuronal excitability of LEC L5 pyramidal neurons requires recall of engram specific cues with associated novel information. **a,** *Top*: Schematic and behavioural data for *ex vivo* recordings from mice which were not presented with a second test phase before slicing, bars represent mean ± SEM (n = 10 mice). *Bottom:* action potential firing (mean ± SEM), representative voltage responses to −100 and +200 pA current injections for a tdTom^-^ and tdTom^+^ cell pair, average action potential waveforms, input resistance and afterhyperpolarisation (AHP) amplitude of LEC L5 pyramidal neurons. Boxplots show median (centre line), 25^th^ and 75^th^ percentiles, whiskers show minimum and maximum points. tdTom^-^ n = 21 cells from 10 mice, tdTom^+^ n = 20 cells from 10 mice. **b.** Same as (a) but mice shown familiar object configuration before slicing. tdTom^-^ n = 31 cells from 7 mice, tdTom^+^ n = 30 cells from 7 mice. **c**. Same as (a) but mice shown a novel object set before slicing. tdTom^-^ n = 31 cells from 9 mice, tdTom^+^ n = 32 cells from 9 mice. Scale bars 200 ms, 40 mV. * = p<0.05. See Supplementary Data Tables 6-8 for further parameters and statistical analysis.

**Table 5.** Summary of physiological properties of layer 5 LEC tdTom- and tdTom+ neurons recorded from ex vivo slices from animals which did not undergo a second test phase (related to Fig. 4a). Values represent median (25,75 percentiles). Reported statistical values show Mann Whitney U test with 2-tailed p-value, except for ADP presence which uses Chi-squared test with 2-tailed p value. RMP resting membrane potential, R_input_ input resistance, AP action potential, AHP afterspike-hyperpolarisation, mAHP medium AHP, ADP afterspike-depolarisation, RoR rate of rise

| Physiological property | tdTom- L5 LEC pyramidal cells | tdTom+ L5 LEC pyramidal cells | Mann Whitney U | P-value |
| --- | --- | --- | --- | --- |
| n (cells, mice) | 21, 10 | 20, 10 |  |  |
| n cells (DLE, DIE, VIE) | 19, 1, 1 | 18, 1, 1 |  |  |
| RMP (mV) | -62.8 (-65.0,-59.5) | -62.3 (-63.6,-58.0) | 173.5 | 0.348 |
| $R_{input}$ (MΩ) | 82.6 (68.6,127.0) | 71.7 (53.3,111.6) | 172 | 0.332 |
| Tau (ms) | 15.8 (12.9,21.2) | 16.2 (13.6,19.4) | 209 | 0.990 |
| Sag (%) | 12.6 (6.1,15.9) | 10.9 (2.7,24.6) | 201 | 0.826 |
| Rheobase (pA) | 126.7 (101.5,165.5) | 152.7 (88.1,181.8) | 190 | 0.615 |
| AP threshold (mV) | -36.5 (-40.7,-32.5) | -38.4 (-42.9,-34.6) | 158 | 0.181 |
| AHP (mV) | -5.9 (-11.0,-4.1) | -5.6 (-8.8,-3.3) | 180 | 0.446 |
| mAHP amplitude (mV) | -1.8 ( -2.9,-0.6) | -1.5 (-3.1,-0.5) | 207 | 0.949 |
| mAHP area (mV.ms) | -0.13 (-0.23,-0.03) | -0.11 (-0.24,-0.03) | 203 | 0.867 |
| ADP presence (#cells) | 4/21 | 5/20 | $\chi^2 = 0.21$ | 0.645 |
| ADP amplitude (mV) | 0.60 (0.11,2.25) | 0.35 (0.16,1.45) | 8 | 0.730 |
| AP amplitude (mV) | 63.3 (59.7,69.2) | 73.3 (62.6,80.7) | 146 | 0.098 |
| AP half width (ms) | 0.92 (0.77,0.97) | 0.84 (0.79,0.94) | 195 | 0.703 |
| AP max RoR (mV.ms<sup>-1</sup>) | 241 (212,322) | 294 (252,381) | 148 | 0.109 |
| **Cell firing** | **Mixed-effects model** | **Factor** | **F** | **P-value** |
| | | Current | $F_{(1.6, 60.35)} = 153$ | $<1 \times 10^{-15}$ |
| | | Labelling | $F_{(1,39)} = 0.003$ | 0.96 |
| | | Interaction | $F_{(4,156)} = 0.55$ | 0.76 |

OiP recall in our experiments contains both static and displaced objects, thus the increased excitability of LEC neurons may result from either memory recall or encoding of new information. To determine whether excitability changes could be induced only by recall, mice were re-exposed to the familiar object configuration (previously encountered in test phase 1) immediately prior to slicing (Fig. 4B). Re-exposure to this familiar object configuration did not, however, lead to any changes in firing or intrinsic electrophysiological properties (Fig. 4B, Table 6), suggesting that retrieval of specific cues in a familiar configuration does not alter engram cell excitability.

**Table 6.** Summary of physiological properties of layer 5 LEC tdTom- and tdTom+ neurons recorded from ex vivo slices from animals exposed to familiar object configuration at test 2 (related to Fig. 4b). Values represent median (25,75 percentiles). Reported statistical values show Mann Whitney U test with 2-tailed p-value, except for ADP presence which uses Chi-squared test with 2-tailed p value. ^a^*data not collected for one cell*. RMP resting membrane potential, R_input_ input resistance, AP action potential, AHP afterspike-hyperpolarisation, mAHP medium AHP, ADP afterspike-depolarisation, RoR rate of rise.

| Physiological property | tdTom- L5 LEC pyramidal cells | tdTom+ L5 LEC pyramidal cells | Mann Whitney U | P-value |
| --- | --- | --- | --- | --- |
| n (cells, mice) | 31, 7 | 30, 7 |  |  |
| n cells (DLE, DIE, VIE) | 29, 2, 0 | 28, 2, 0 |  |  |
| RMP (mV) | -65.3 (-67.9,-62.1) | -64.3 (-66.8,-62.7) | 425.5 | 0.574 |
| $R_{input}$ (MΩ) | 76.1 (56.0,102.3) | 82.0 (56.3,99.6) | 449 | 0.824 |
| Tau (ms) | 17.0 (13.6,19.6) | 15.1 (12.2,19.6) | 434 | 0.662 |
| Sag (%) | 11.7 (6.0,16.9) | 10.3 (5.5,16.8) | 454 | 0.881 |
| Rheobase (pA) | 144.2 (103.6,233.5)<sup>a</sup> | 150.3 (90.40,207.8) | 442 | 0.912 |
| AP threshold (mV) | -39.4 (-41.8,-34.4) | -37.1 (-39.9,-33.6) | 374 | 0.193 |
| AHP (mV) | -7.3 (-10.8,-4.3) | -6.5 (-11.4,-5.1) | 454 | 0.881 |
| mAHP amplitude (mV) | -1.4 (-2.2,-0.8)<sup>a</sup> | -1.8 (-3.0,-0.8)<sup>a</sup> | 389 | 0.493 |
| mAHP area (mV.ms) | -0.11 (-0.17,-0.06)<sup>a</sup> | -0.12 (-0.23,-0.05)<sup>a</sup> | 384 | 0.447 |
| ADP presence (#cells) | 4/30<sup>a</sup> | 8/30 | $\chi^2 = 1.67$ | 0.197 |
| ADP amplitude (mV) | 0.12 (0.09,1.58)<sup>a</sup> | 1.46 (0.45,1.86) | 10 | 0.368 |
| AP amplitude (mV) | 72.0 (63.7,79.4) | 67.1 (60.7,78.5) | 369 | 0.170 |
| AP half width (ms) | 0.81 (0.78,0.91) | 0.84 (0.75,0.90) | 438 | 0.704 |
| AP max RoR (mV.ms<sup>-1</sup>) | 304 (263,377) | 269 (226,362) | 388 | 0.272 |
| **Cell firing** | **Mixed-effects model** | **Factor** | **F** | **P-value** |
| | | Current | $F_{(1.3, 74.68)} = 156$ | $<1 \times 10^{-15}$ |
| | | Labelling | $F_{(1,59)} = 0.004$ | 0.85 |
| | | Interaction | $F_{(4,236)} = 0.34$ | 0.86 |

These data suggest that changes in intrinsic excitability may be a mechanism by which novel information regarding object locations is integrated into the pre-existing engram as part of a reconsolidation-like process. Alternatively, the increased excitability in tdTom neurons may have been a response to a more general novelty signal created by object reconfiguration. To determine this, we presented mice with a previously unencountered set of objects at test phase 2 (Fig. 4C). No difference in firing rate or other measures of cellular excitability were observed in *ex vivo* slices from mice exposed to novel objects, however action potential dynamics showed similar changes (slower rate of rise, smaller amplitude) to those observed in slices from mice which had encountered rearranged engram-related objects (Fig. 4C, Table 7). These data demonstrate that whilst a novel object set can alter action potential dynamics, the increased cellular excitability and firing rates observed in response to object reconfiguration are engram-specific and are not a result of a general novelty signal.

**Table 7.** Summary of physiological properties of layer 5 LEC tdTom- and tdTom+ neurons recorded from ex vivo slices from animals were exposed to a new object set prior to slice preparation (related to Fig. 4c). Values represent median (25,75 percentiles). Reported statistical values show Mann Whitney U test with 2-tailed p-value, except for ADP presence which uses Chi-squared test with 2-tailed p value. RMP resting membrane potential, R_input_ input resistance, AP action potential, AHP afterspike-hyperpolarisation, mAHP medium AHP, ADP afterspike-depolarisation, RoR rate of rise.

| Physiological property | tdTom- L5 LEC pyramidal cells | tdTom+ L5 LEC pyramidal cells | Mann Whitney U | P-value |
| --- | --- | --- | --- | --- |
| n (cells, mice) | 31, 9 | 32, 9 |  |  |
| n cells (DLE, DIE, VIE) | 29, 2, 0 | 29, 3, 0 |  |  |
| RMP (mV) | -63.2 (-64.9,-61.0) | -63.2 (-67.2,-58.1) | 249.5 | 0.997 |
| $R_{input}$ (MΩ) | 81.6 (62.6,102.8) | 88.2 (57.9,125.2) | 445 | 0.490 |
| Tau (ms) | 16.2 (12.7,18.3) | 16.1 (13.8,18.0) | 473 | 0.759 |
| Sag (%) | 14.6 (7.1,21.9) | 11.86 (5.67, 17.77) | 396 | 0.173 |
| Rheobase (pA) | 155.5 (120.1, 236.8) | 156.3 (115.2,199.4) | 442 | 0.465 |
| AP threshold (mV) | -38.0 (-40.5,-35.8) | -36.4 (-39.9,-34.3) | 378 | 0.107 |
| AHP (mV) | -7.9 (-10.2,-4.5) | -6.3 (-8.1,-3.1) | 391 | 0.152 |
| mAHP amplitude (mV) | -2.3 (-3.7,-1.3) | -1.5 (-2.8,-0.8) | 378 | 0.107 |
| mAHP area (mV.ms) | -0.16 (-0.27,-0.10) | -0.12 (-0.21,-0.06) | 389 | 0.144 |
| ADP presence (#cells) | 16/31 | 9/32 | $\chi^2 = 1.91$ | 0.057 |
| ADP amplitude (mV) | 0.48 (0.17,0.88) | 0.38 (0.24,1.38) | 71 | 0.978 |
| AP amplitude (mV) | 71.3 (63.4,78.4) | 64.2 (55.4,73.4) | 337 | 0.029 |
| AP half width (ms) | 0.79 (0.72,0.87) | 0.88 (0.79,0.94) | 289 | 0.004 |
| AP max RoR (mV.ms<sup>-1</sup>) | 293 (240,373) | 242 (200,302) | 298.5 | 0.006 |
| **Cell firing** | **Mixed-effects model** | **Factor** | **F** | **P-value** |
| | | Current | $F_{(1.3, 81.67)} = 182$ | $<1 \times 10^{-15}$ |
| | | Labelling | $F_{(1,61)} = 0.001$ | 0.97 |
| | | Interaction | $F_{(4,244)} = 0.23$ | 0.92 |

## Discussion

In this study we investigate the identity of cortical neurons encoding associative recognition memories, showing that engram cell recruitment and reactivation upon memory retrieval occurs across the lateral entorhinal, medial prefrontal and perirhinal cortices. However, within these regions, only the dorsal extent of the LEC showed a strong correlation between engram-cell reactivation and memory performance, and reactivation was higher in L5/6 DLent neurons. Furthermore, reactivation of LEC engram neurons was shown to be necessary for memory recall, and layer 5 LEC neurons exhibited enhanced intrinsic excitability following a memory-recall task. Crucially, increased excitability was contingent upon the object set having been rearranged at recall whereas unmoved objects or a second set of objects did not increase excitability. These data collectively implicate deep layer DLent neurons as crucial for object-place associations, and increased excitability of these neurons may represent a novel mechanism by which new information is integrated into an existing memory engram.

Recent studies using molecular-genetic activity-labelling have provided considerable support to the notion of memory engrams as cellular substrates of memory (Josselyn and Tonegawa, 2020; Ortega-de San Luis et al., 2023), however these investigations have relied on strongly aversive or rewarding behaviours. Here we provide a detailed characterisation of cortical engrams for associative recognition memory, a spontaneous, unrewarded, single-trial form of learning which is critical to our everyday lives.

Our analysis showed that OiP produces robust activity-dependent labelling of neurons in multiple recognition memory associated cortices. However, memory performance correlated with reactivation of tdTom cells only in LEC, and this correlation exhibited a dorsal-ventral gradient, with a strong correlation in DLent and no correlation in VIent. These data support the notion that DLent can support object-place associations, in contrast with PRH, which is thought to only encode object identity and novelty (Deshmukh et al., 2012) and in which no correlation was found between reactivation and memory.

Within LEC, L5/6 neurons showed higher levels of reactivation than L2/3 neurons, despite similar numbers of neurons in these regions being activated overall at encoding and recall; this heightened level of reactivation combined with the intrinsic excitability plasticity seen in these neurons implicates DLent L5/6 as a key locus of object-place associations. Whilst previous studies have shown object-place representations in LEC L5/6 (Keene et al., 2016), most evidence to date has implicated superficial LEC neurons in encoding associative (Lee et al., 2021) and episodic-like memory (Vandrey et al., 2020). L5 LEC neurons are anatomically well positioned within associative recognition memory circuitry, being the recipients of convergent top-down feedback inputs from both dorsal hippocampus(Ohara et al., 2023) and inputs from mPFC (Kinnavane and Banks, 2022; Kinnavane et al., 2025). Outputs from L5 LEC neurons project to other telencephalic regions (Ohara et al., 2023), including mPFC (Hoover and Vertes, 2007), and locally within LEC where they target superficial layers (Ohara et al., 2021) which project to hippocampus. Dorsal LEC L5 is, therefore, a key node in bidirectional loops between LEC-hippocampus and LEC-mPFC.

The notion that deep-layer LEC neurons support object-place associations is supported by plasticity of excitability which occurred specifically in LEC L5 TRAP2 pyramidal neurons following exploration of reconfigured objects, which was absent in L2a fan cells, PRH L5, and both L2/3 and L5 mPFC pyramidal neurons. This finding raises the possibility that plasticity of cellular excitability in these neurons may represent a mechanism by which neural object-place representations can be rapidly altered in response to engram-related environmental novelty. Whilst changes in cellular excitability were only found in LEC L5, this is not to say that other plastic changes, such as synaptic plasticity, do not occur in other cell populations (Ryan et al., 2015; Lee et al., 2023; Tozzi et al., 2024).

Two alterations to LEC L5 tdTom^+^ cells were found in this study: firstly, increased input resistance, with an accompanying increase in firing upon depolarisation. A decrease in potassium conductance is a potential mechanism for this increased excitability associated with recall (Pignatelli et al., 2019), whereas other mechanisms regulating firing rate such as AHP (Disterhoft and Oh, 2006; Cai et al., 2016) were unaltered. Secondly, we observed action potentials had reduced amplitude and rate of rise, likely reflecting decreased voltage-gated sodium channel function which has previously been observed in response to dopaminergic activity (Cantrell and Catterall, 2001), which signals novelty in LEC (Lee et al., 2021).

Increased engram cell excitability was only observed under specific conditions. The absence of plasticity in tdTom^+^ cells from mice which did not re-encounter objects before slicing shows that increased excitability either leading to, or resulting from, recruitment to an engram (Disterhoft and Oh, 2006; Josselyn and Frankland, 2018) either did not occur or was not persistent 48 h later. Furthermore, as we found no electrophysiological difference between engram and non-engram cells when no recall had taken place before slicing, it is unlikely that engram recruitment is biased towards a particular subclass of LEC L5 pyramidal cells, which have different electrophysiological phenotypes, such as layer 5a vs 5b pyramidal cells (Ohara et al., 2021).

Re-exposure to objects in familiar locations did not alter engram cell excitability. Meanwhile, exploration of completely novel objects resulted only in altered action potential dynamics. We hypothesize that a novelty signal, such as release of neuromodulatory transmitter, may be sufficient to induce changes in action potential dynamics in both the novel objects and reconfigured objects groups, but that this novelty signal must be accompanied by strong activation of synapses within the engram network, including from engram cells in neurons from distal regions such as mPFC and hippocampus, to generate a plasticity signal that results in increased firing.

The function of intrinsic plasticity in engram cells is unclear. Evidence suggests that engram cell intrinsic excitability increases during a short temporal window after memory recall, and that these changes facilitate pattern separation and completion (Pignatelli et al., 2019). Heightened excitability may therefore act as a (likely temporary) mechanism of updating environmental representations by integrating new information into an established engram via a re-consolidation like process, and may facilitate processes underlying systems consolidation (such as replay events (Olafsdottir et al., 2018)) which facilitate longer term storage of environmental changes in the form of synaptic changes (Ryan et al., 2015; Lee et al., 2023).

## Additional resources

Data and code used in this manuscript are available upon request.

## Author Contributions

Research was designed by: P.J.B., G.R.I.B., E.C.W. and Z.I.B. Research was performed by: P.J.B., G.R.I.B., C.A.B. and L.K. Data analysis was performed by: P.J.B., G.R.I.B. and C.A.B. The first draft of the paper was written by P.J.B., review and editing of the paper was performed by: P.J.B., G.R.I.B., C.A.B., E.C.W. and Z.I.B.

## Acknowledgements

This work was supported by a Wellcome Trust Joint Investigator Award to E.C.W. and Z.I.B. (206401/Z/17/Z), a BBSRC project grant to Z.I.B., E.C.W and P.J.B. (BB/X000915/1) and a BBSRC project grant to E.C.W and G.R.I.B. (BB/Y006402/1). We thank Dr Emma Cahill for helpful comments on the manuscript.

---

**Supplementary Figure 1.**
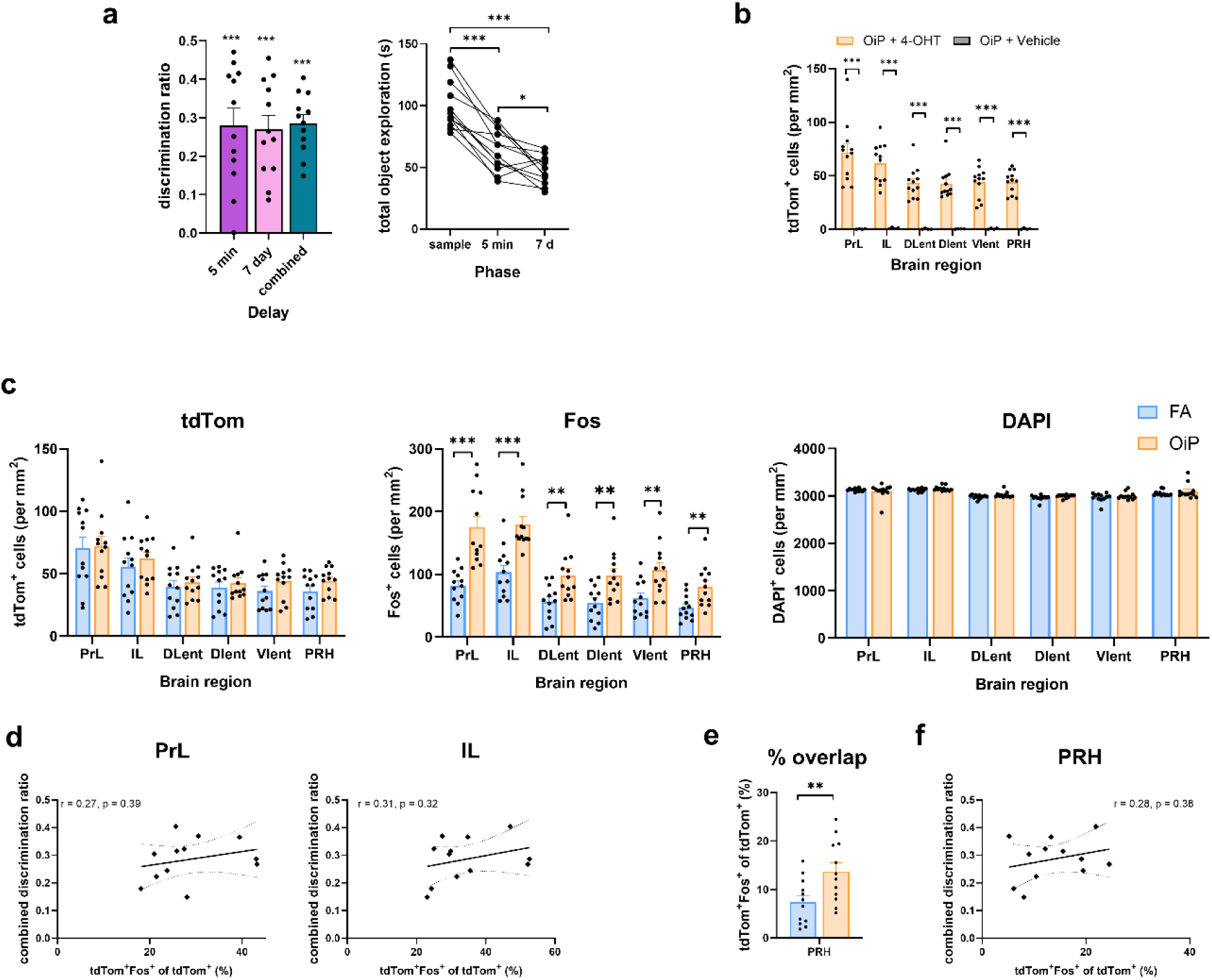
Behavioural and labelling data from FA and OiP tasks. **a**: (Left) Animals showed significant discrimination between the moved and unmoved objects in the OiP task, as measured by the discrimination ratio, at 5 min and 7 d delays and when combined (5 min t_(11)_ = 7.55, p = 1.15 x10^-5^; 7 day t_(11)_ = 6.31, p = 5.79 x 10^-5^; combined t_(11)_ = 12.79, p = 6.04 x 10^-8^). (Right) Total object exploration significantly decreased across each phase of the behavioural task (F_(2,22)_ = 46.99, p = 1.14 x 10^-8^, post-hoc sample vs 5 min p = 2.82 x 10^-5^, sample vs 7 day: p = 1.91 x 10^-5^, 5 min vs 7 day: p = 0.032). **b**: Expression of tdTom was dependent on injection of 4-OHT in all brain regions (in each brain region one-way ANOVA F_(1,14)_ > 28, p < 0.001). OiP + 4-OHT n = 12, OiP + vehicle n = 4. **c**: tdTom, Fos and DAPI cell counts in subregions of mPFC and LEC and in PRH for FA and OiP conditions. **d**: No significant correlation between tdTom and Fos overlap and memory performance in either PrL or IL subregions of mPFC. Dashed line show 95% confidence intervals. **e**: Overlap between tdTom and Fos expression in the perirhinal cortex is significantly higher in the OiP compared to the FA condition. **f**: No significant correlation between tdTom and Fos overlap and memory performance in the perirhinal cortex. Dashed line show 95% confidence intervals. All bar charts show mean ± SEM. * p < 0.05, ** p < 0.01, *** p < 0.001. For FA vs OiP comparisons n = 12 animals for each condition. See Supplementary Data Table 1 for detailed statistical analysis.

**Supplementary Figure 2.**
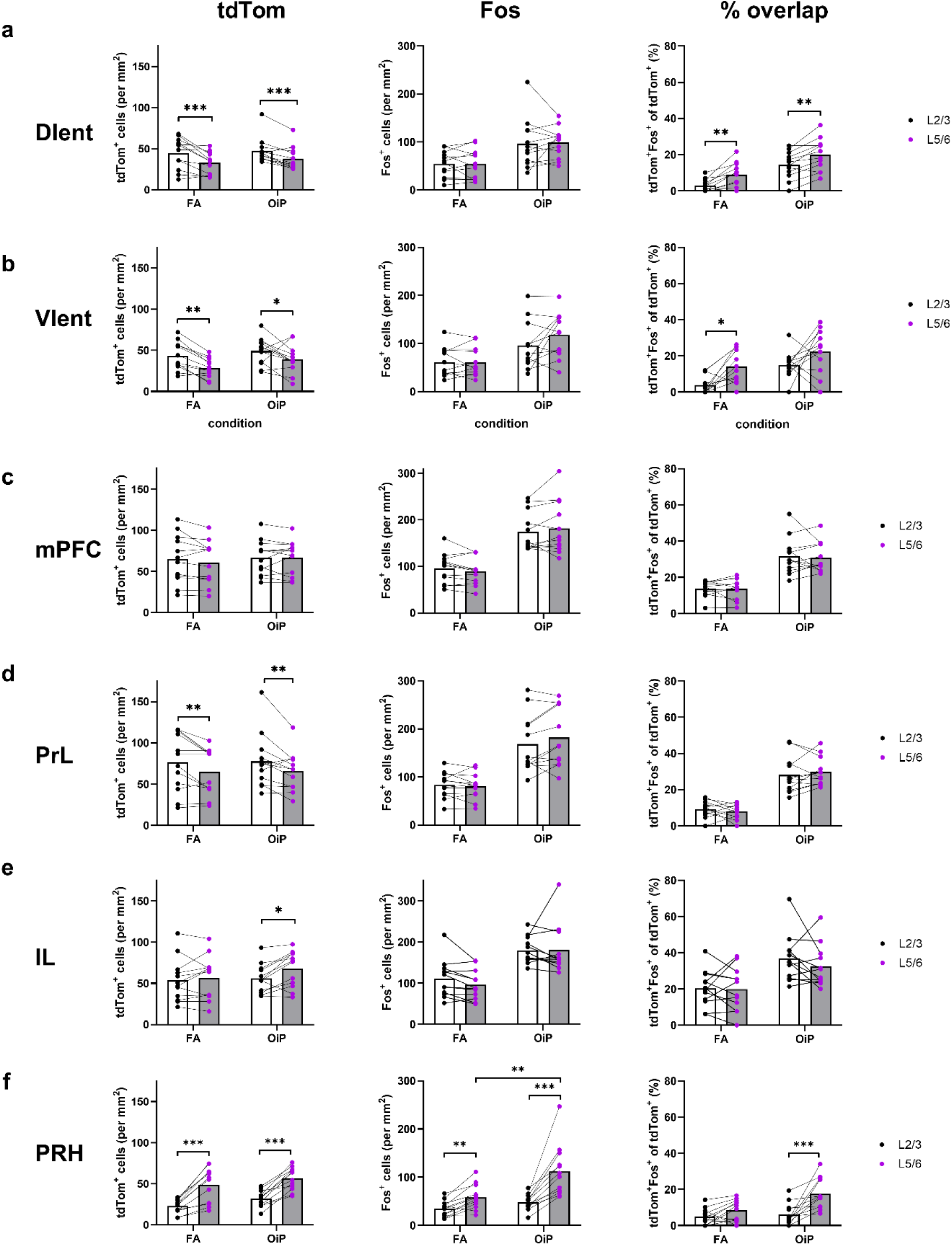
Expression patterns of tdTom, Fos and overlap in layer 2/3 and layer 5/6 across cortical regions. **a**: (Left to right) tdTom, Fos and percent overlap for L2/3 and L5/6 in DIent. **b**: Same as (**a**) but for VIent. **c**: Same as (**a**) but for mPFC. **d**: Same as (**a**) but for PrL. **e**:Same as (**a**) but for IL. **f**: Same as (**a**) but for PRH. Bars represent mean values. *** p < 0.001, ** p < 0.01, * p < 0.05. n = 12 mice per condition. See Supplementary Data Table 2 for detailed statistical analysis.

**Supplementary Data Table 1:** Statistical analysis of tdTom and Fos expression and % overlap across mPFC and LEC (and subdivisions) and PRH. Data shows outcome of one-way ANOVA with behavioural task as a between subjects factor. For both conditions n = 12 mice.

| Brain region | tdTom +ve cells |  | Fos +ve cells |  | % Overlap |  |
| --- | --- | --- | --- | --- | --- | --- |
| | $F_{(1,22)}$ | P-value | $F_{(1,22)}$ | P-value | $F_{(1,22)}$ | P-value |
| **mPFC** | 0.17 | 0.686 | 26.90 | $3.35 \times 10^{-5}$ | 38.54 | $3.01 \times 10^{-6}$ |
| PrL | 0.01 | 0.909 | 24.88 | $5.42 \times 10^{-5}$ | 58.95 | $1.16 \times 10^{-7}$ |
| IL | 0.55 | 0.466 | 21.20 | $1.38 \times 10^{-4}$ | 11.78 | 0.002 |
| **LEC** | 0.89 | 0.356 | 10.33 | 0.004 | 37.42 | $3.71 \times 10^{-6}$ |
| DLent | 0.40 | 0.534 | 9.40 | 0.006 | 26.26 | $3.90 \times 10^{-5}$ |
| Dlent | 0.39 | 0.537 | 9.83 | 0.005 | 20.59 | $1.62 \times 10^{-4}$ |
| Vlent | 2.01 | 0.171 | 9.33 | 0.006 | 27.62 | $2.85 \times 10^{-5}$ |
| **PRH** | 2.60 | 0.121 | 9.26 | 0.006 | 7.64 | 0.011 |

**Supplementary Data Table 2:**
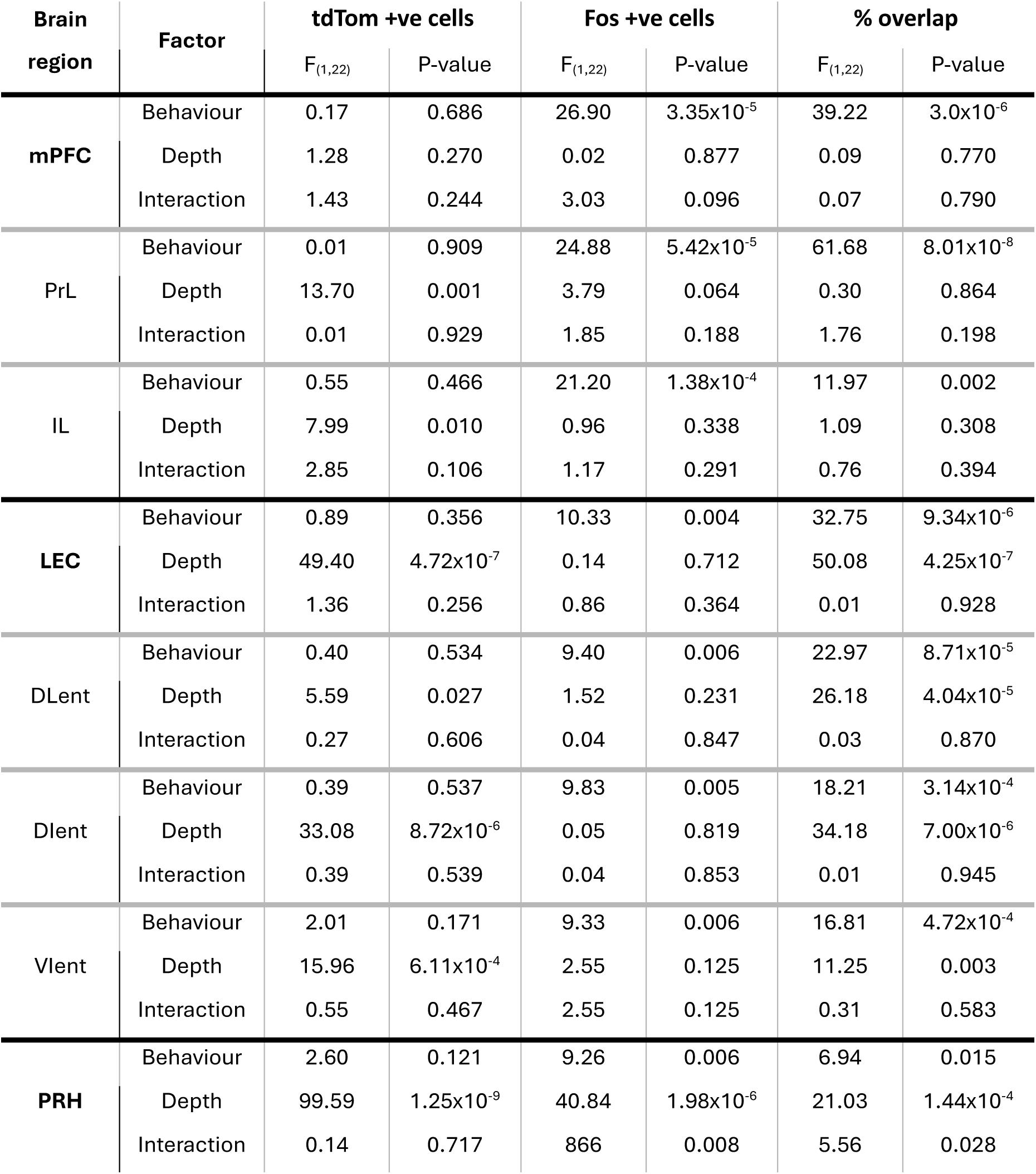
Statistical analysis of tdTom and Fos expression and % overlap in superficial and deep layers of PFC, LEC and PRH. Data shows outcome of two-way ANOVA with behavioural condition (beh. cond.) as a between-subjects factor and depth as a within-subjects factor. n = 12 mice per condition.

